# NEDD4-binding protein 1 suppresses HBV replication by degrading HBV RNAs

**DOI:** 10.1101/2023.10.06.561254

**Authors:** Nobuhiro Kobayashi, Saori Suzuki, Yuki Sakamoto, Rigel Suzuki, Kento Mori, Yume Kosugi, Lihan Liang, Tomoya Saito, Takuma Izumi, Kisho Noda, Daisuke Okuzaki, Yumi Kanegae, Sanae Hayashi, Yasuhito Tanaka, Yoshiharu Matsuura, Osamu Takeuchi, Tomokazu Tamura, Akinobu Taketomi, Takasuke Fukuhara

**Author notes:** These authors contributed equally (Nobuhiro Kobayashi, Saori Suzuki). Address correspondence to Fukuhara Takasuke,. Author order was determined by the basis of contribution (see Acknowledgement for detail). **Conflict of Interest statement** Nothing to report.

## Abstract

Chronic infection with hepatitis B virus (HBV) places patients at increased risk for liver cirrhosis and hepatocellular carcinoma. Although nucleos(t)ide analogs are mainly used for the treatment of HBV, they require long-term administration and may lead to the emergence of drug resistant mutants. Therefore, to identify targets for the development of novel anti-HBV drugs, we screened for HBV-suppressive host factors using a plasmid expression library of RNA-binding proteins (RBPs). We screened 132 RBPs using an expression plasmid library by measuring HBV relaxed circular DNA (rcDNA) levels in hepatocellular carcinoma. Our screen identified NEDD4-binding protein 1 (N4BP1) as having an anti-HBV effect. In hepatocellular carcinoma cell lines transfected or infected with HBV, overexpression of N4BP1 decreased rcDNA levels while knockdown or knockout of the gene encoding N4BP1 rescued rcDNA levels. N4BP1 possesses the KH-like and RNase domains and both were required for the anti-HBV effect of N4BP1. Additionally, we measured levels of HBV pregenomic RNA (pgRNA) and covalently closed circular DNA (cccDNA) in the RBP-transfected cells and confirmed that N4BP1 binds pgRNA directly and degraded both the 3.5 kb and 2.4/2.1 kb HBV RNA. In summary, N4BP1 is a newly identified host factor able to counteract HBV production by promoting the degradation of 3.5 kb and 2.1/2.4 kb HBV RNA.

**Importance:** There is still a large number of HBV-infected people in the world today because of no curative treatment for HBV infection. In this study, we focused on and screened RNA-binding proteins to identify new host factors which inhibit HBV replication. As a result, we found that NEDD4-binding protein 1 (N4BP1) expression suppresses rcDNA production by promoting the degradation of pregenomic RNA, 2.4kb and 2.1kb HBV RNA. Furthermore, KH-like domain or RNase domain of N4BP1 were involved in this anti-HBV effect. In addition, the N4BP1 levels were lower in HCC resection samples of exacerbated patients, suggesting that individual N4BP1 levels might be related to HCC progression. This novel factor can potentially become a key to new HBV treatments.

## Introduction

Hepatitis B virus (HBV) infection causes either acute or chronic hepatitis, with approximately 296 million people worldwide suffering from chronic hepatitis B and an estimated 1.5 million new cases arising each year. Chronically infected individuals are at increased risk for liver disease, and annually ∼820,000 die from cirrhosis and hepatocellular carcinoma (HCC) caused by HBV infection (1). Although there is a prophylactic vaccine for HBV, there are still many new cases of HBV infection due to the large number of chronic HBV infected patients still in Africa, Asia, and Eastern Europe and the low vaccination rates in these highly endemic areas (2).

HBV is an enveloped DNA virus with a small, partially double-stranded genome of 3.2 kbp. Entry into hepatocytes is mediated by binding to the bile acid transporter sodium-taurocholate cotransporting polypeptide (NTCP). After being released into the nucleus, the HBV genome is converted from its relaxed circular DNA (rcDNA) form to covalently closed circular DNA (cccDNA) using the host DNA repair system (3-6). The cccDNA serves as the transcription template for viral mRNAs, including the HBV pregenomic RNA (pgRNA). pgRNA is used to synthesize the negative-strand DNA first and then the positive-strand is synthesized to generate rcDNA.

Interferon (IFN) and nucleos(t)ide analogs (NAs) are currently used as treatments for HBV infection, and both have shown efficacy in controlling viremia (7, 8). IFNs are involved in the inhibition of HBV replication, transcription, and other important processes by inducing the expression of various IFN-stimulated genes (ISGs) in host cells (9). However, IFN treatment can cause considerable side effects, including liver damage, and is thus unsuitable for patients with severe liver dysfunction (10).

In contrast, NAs impair further HBV intrahepatic spreading by blocking production of new virions. But the HBV antigens HBs or HBe are not eliminated with treatment as NAs inhibit the reverse transcription step that occurs after these antigens have already been translated. Furthermore, NAs do not directly target cccDNA, allowing the virus to persist in hepatocytes (11, 12). Consequently, to prevent relapse due to reactivation of HBV after discontinuing NAs, long-term treatment is recommended (13).

In efforts to overcome the limitations of these current therapeutic options, research has looked to identify host factors with an antiviral role. Understanding the mechanism of action of such factors could aid in the development of new antiviral treatments. Several host factors are known to inhibit HBV replication, including RNA-binding proteins (RBPs). For example, the CCCH zinc finger antiviral protein ZAP (ZC3HAV1) binds to HBV RNA with its four zinc finger motifs at the N-terminus, promoting pgRNA degradation (14). Similarly, MCPIP1 (ZC3H12A, regnase-1) has also been reported to inhibit HBV replication by binding to HBV pgRNA through its zinc finger motif, directly degrading pgRNA via its RNase activity (15). Furthermore, PUF60 is a host factor that not only positively regulates HBV 3.5 kb precore plus pregenomic RNA expression by binding to the HBV enhancer region II with the transcription factor TCF7L2 but can also promote degradation of the HBV 3.5 kb RNA by binding via its N-terminal RNA recognition motif (16). Despite these descriptions, HBV produces high levels of HBV RNAs in infected primary human hepatocytes (PHHs) and i*n vivo*, so the antiviral effects of these host factors appear limited.

To determine if other RBPs may have an antiviral role, we performed a comprehensive screen using an expression library of RBPs. We identified NEDD4-binding protein 1 (N4BP1) as a host factor with anti-HBV activity. We subsequently elucidated the mechanism of how N4BP1 suppresses HBV replication by promoting the degradation of HBV RNAs.

## Results

### Screen for RNA-binding proteins that inhibit HBV propagation

To identify novel RBPs involved in HBV propagation, we screened 132 RBPs using an expression plasmid library. Huh7 cells were co-transfected with an RBP expression plasmid and a 1.3-fold-overlength HBV genome (genotype C). At 3 days post-transfection (d.p.t.), cells were collected to measure intracellular rcDNA in core particles. As shown in Fig 1A, 23 out of 132 RBPs suppressed rcDNA levels by at least 50% compared to the control.

**Fig 1.**
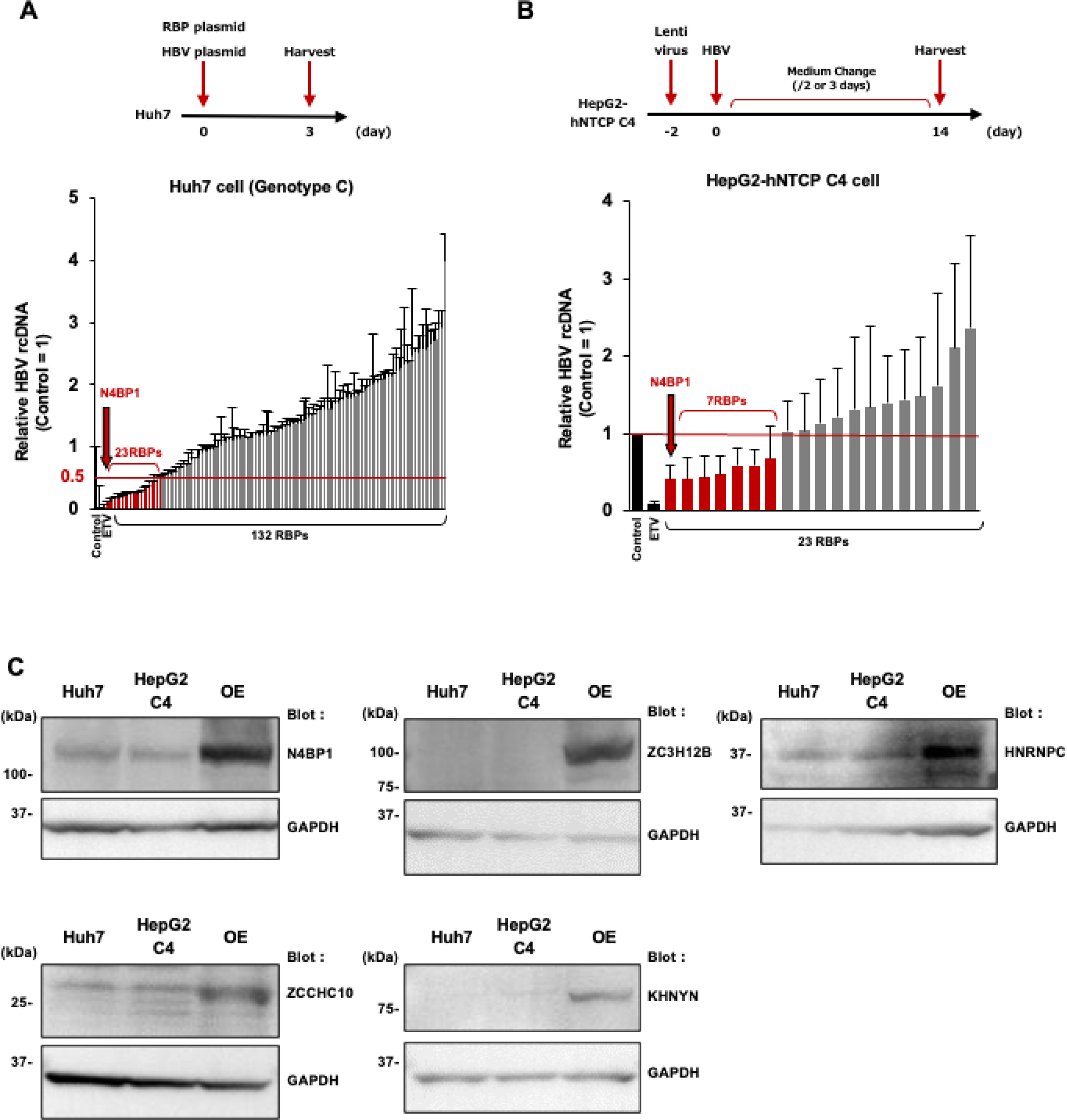
Screening for RNA binding proteins (RBPs) that inhibit HBV propagation. (A) A plasmid containing the 1.3-fold-overlength genome of HBV genotype C and 132 RBP expression plasmids were co-transfected into Huh7 cells. Intracellular rcDNA levels were assessed at 3 d.p.t.. Using the 50% of rcDNA level of the control as a cutoff, 23 out of the 132 RBPs suppressed HBV rcDNA levels. (B) HepG2-hNTCP C4 cells overexpressing the 23 RBPs were infected with HBV genotype D. At 14 d.p.i., cells were harvested and intracellular rcDNA levels were measured by qPCR. Seven out of the 23 RBPs (N4BP1, TIA-1, ZC3H12B, PJA1, HNRNPC, ZCCHC10, and KHNYN) reduced rcDNA levels in comparison to the control. (C) WB analyses in naïve Huh7 cells, HepG2-hNTCP C4 cells of the five factors (N4BP1, ZC3H12B, HNRNPC, ZCCHC10, and KHNYN) identified in the secondary screening. As a positive control, WB were performed on Huh7 cells transfected to overexpress each factor. Error bars represent the SD. Abbreviations: RBP (RNA binding protein), ETV (entecavir), HepG2 C4 (HepG2-hNTCP C4), OE (overexpression)

As a secondary screen, we then examined the effect of these 23 RBPs on HBV entry and replication. HepG2-hNTCP C4 cells expressing each of the 23 RBPs were infected with HBV (genotype D), and intracellular rcDNA was measured by quantitative PCR (qPCR) at 14 days post-infection (d.p.i.). Seven of the 23 RBPs (N4BP1, TIA-1, ZC3H12B, PJA1, HNRNPC, ZCCHC10, and KHNYN) reduced rcDNA levels in comparison with the control (Fig 1B). These results were consistent with previous reports that found TIA-1 and PJA1 inhibit expression of HBs antigen and HBV transcription and replication, respectively (17, 18).

Of the five remaining factors, we found by western blot that three (N4BP1, HNRNPC, and ZCCHC10) are endogenously expressed in Huh7 and HepG2-hNTCP C4 cells (Fig 1C). To test whether these proteins affect HBV replication, we knocked down each one and assessed rcDNA levels following transfection with the 1.3-fold-overlength genome of HBV genotype C. The antiviral drug entecavir, a nucleoside analog, was used as a positive control. Knockdown of each factor was confirmed by western blotting (WB), but unlike knockdown of N4BP1, knockdown of HNRNPC or ZCCHC10 did not consistently increase the levels of rcDNA depending on the siRNA used (Fig 2A-2C). Thus, N4BP1 became the focus of our study.

**Fig 2.**
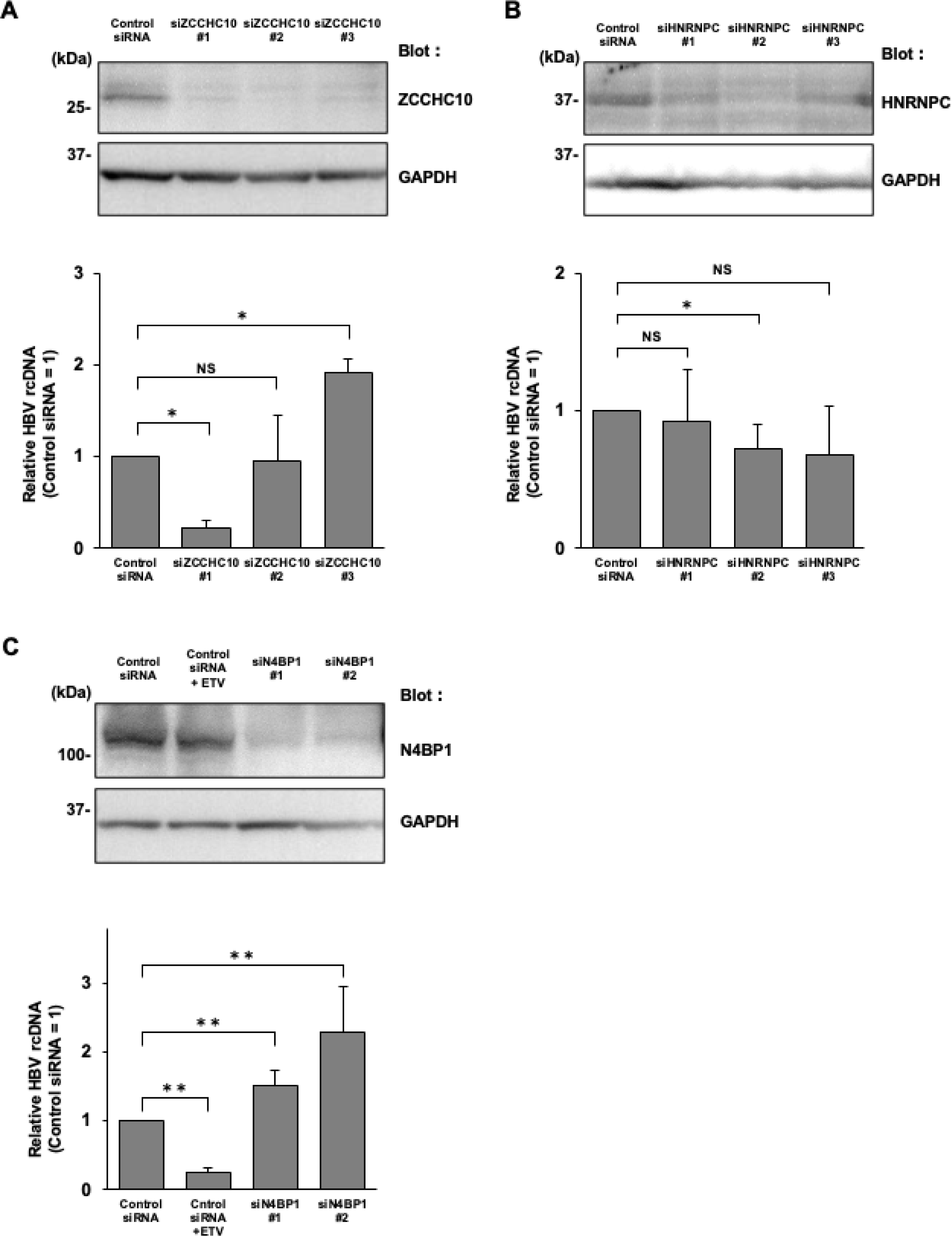
Measurement of rcDNA levels after RBP knockdown. (A) Huh7 cells were transfected with siRNA, then co-transfected with the 1.3-fold-overlength genome of HBV genotype C and an RBP expression plasmid. WB analyses and qPCR of rcDNA were performed at 3 d.p.t.. ZCCHC10 (A), HNRNPC (B), and N4BP1 (C). GAPDH was used as a loading control and ETV was used as a positive control. Error bars represent the SD. *P < 0.05, **P < 0.01, as determined by Mann-Whitney U test. All qPCR experiments were independently performed at least three times. Abbreviations: NS (not significant)

### N4BP1 is a suppressive host factor of HBV replication

Next, to determine whether N4BP1 also had an antiviral effect against other HBV genotypes, we co-transfected Huh7 cells with a plasmid expressing N4BP1 and a plasmid containing 1.3-fold-overlength genomes of HBV genotypes A, B, and C. At 3 d.p.t., rcDNA levels were decreased in cells transfected with each genotype (Fig 3A).

**Fig 3.**
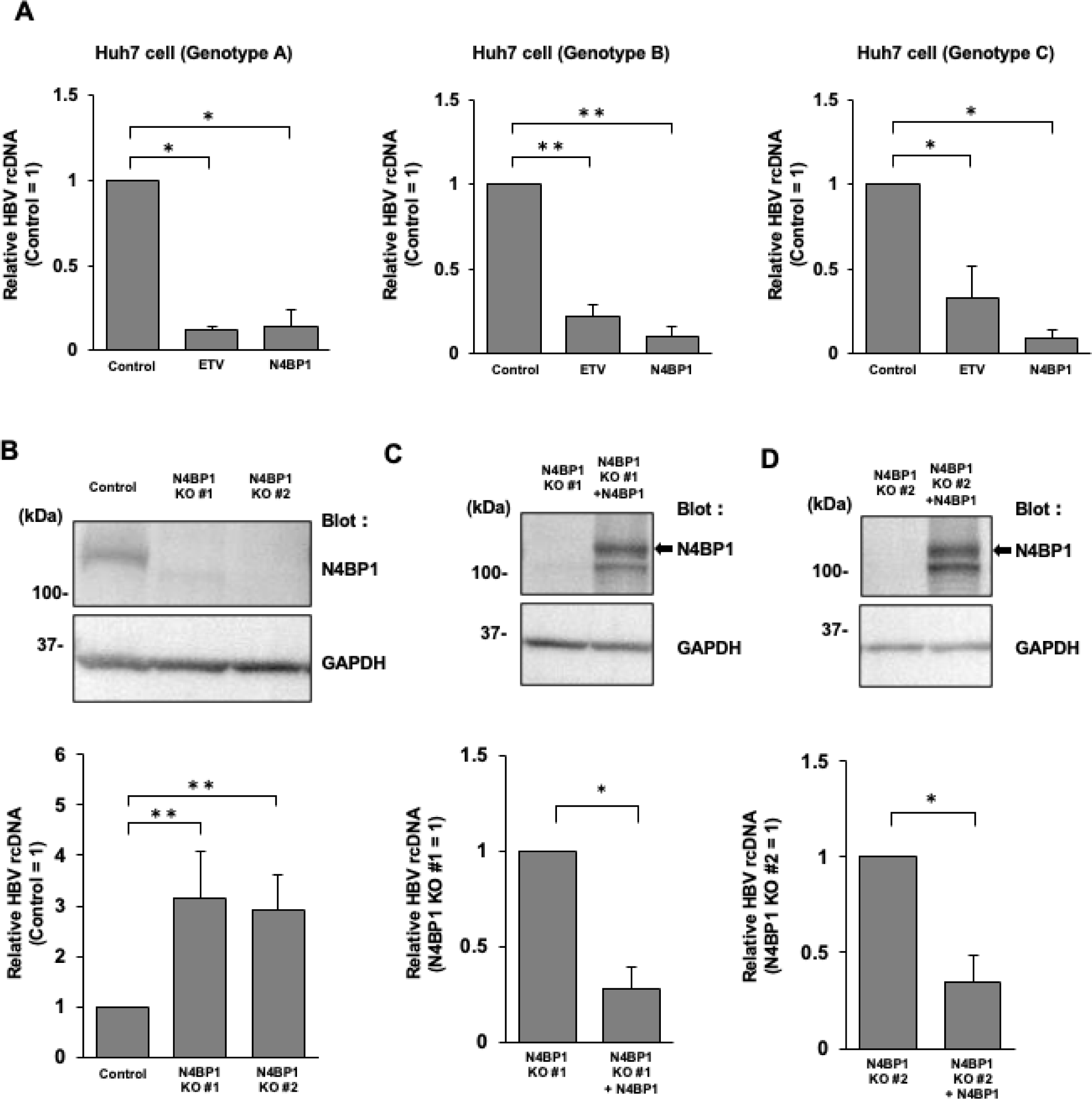
N4BP1 is a suppressive host factor of HBV replication. (A) Plasmids containing 1.3-fold-overlength genomes of HBV genotypes A, B, or C were co-transfected with N4BP1 expression plasmids into Huh7 cells. At 3 d.p.t., the rcDNA levels were measured by qPCR. Genotype A (left), genotype B (middle), and genotype C (right). (B) WB analysis and rcDNA levels of N4BP1 KO Huh7 cells. N4BP1 KO Huh7 cells were transfected with the plasmid containing 1.3-fold-overlength genomes C and WB and qPCR were performed at 3 d.p.t.. N4BP1 expression by WB (upper, anti-N4BP1). GAPDH was used as a loading control (lower, anti-GAPDH). rcDNA levels were increased relative to control in both N4BP1-KO lines that were generated. N4BP1 KO Huh7 cell lines #1 (C) and #2 (D) were transfected with the N4BP1 plasmid to restore N4BP1 expression (upper, anti-N4BP1). GAPDH was used as a loading control (lower, anti-GAPDH). In the N4BP1-rescued KO Huh7 cells, the level of rcDNA was significantly lower than in the non-rescued N4BP1 KO Huh7 cells. Error bars represent the SD. *P < 0.05, **P < 0.01, as determined by Mann-Whitney U test. All qPCR experiments were independently performed at least three times.

To further elucidate the role of N4BP1 in HBV replication, we constructed Huh7 N4BP1 knockout (KO) cells using CRISPR/Cas9. We generated two clones of N4BP1 KO Huh7 cells (Fig 3B) and once more assessed whether intracellular rcDNA levels changed at 3 d.p.t. with a plasmid containing the 1.3-fold-overlength genome of HBV genotype C. In both KO lines, rcDNA levels increased approximately 3-fold compared to control cells (Fig 3B). After rescuing N4BP1 expression in our KO line, rcDNA levels became significantly decreased again (Fig 3C, 3D), suggesting that N4BP1 is a suppressive host factor of HBV replication.

In addition, we performed a transcriptomic analysis of N4BP1-overexpressing PHHs vs control PHHs to assess whether N4BP1 affects regulatory transcription factors that might affect HBV replication (Fig. 4). We set a cutoff value of gene expression at absolute fold change greater than 2 and an adjusted P-value less than 0.1 compared to control cells and found 178 genes upregulated in N4BP1-overexpressing vs control PHHs and 230 genes downregulated (Fig 4A-B, S1 Table). None of these factors were regulatory transcription factors. In addition, pathway enrichment analysis did not show significant coordinated changes in genes associated with translational pathways (Fig 4C). These findings suggest that N4BP1 overexpression was not associated with overarching changes in the expression of regulatory transcription factors that would also affect HBV RNA production.

**Fig 4.**
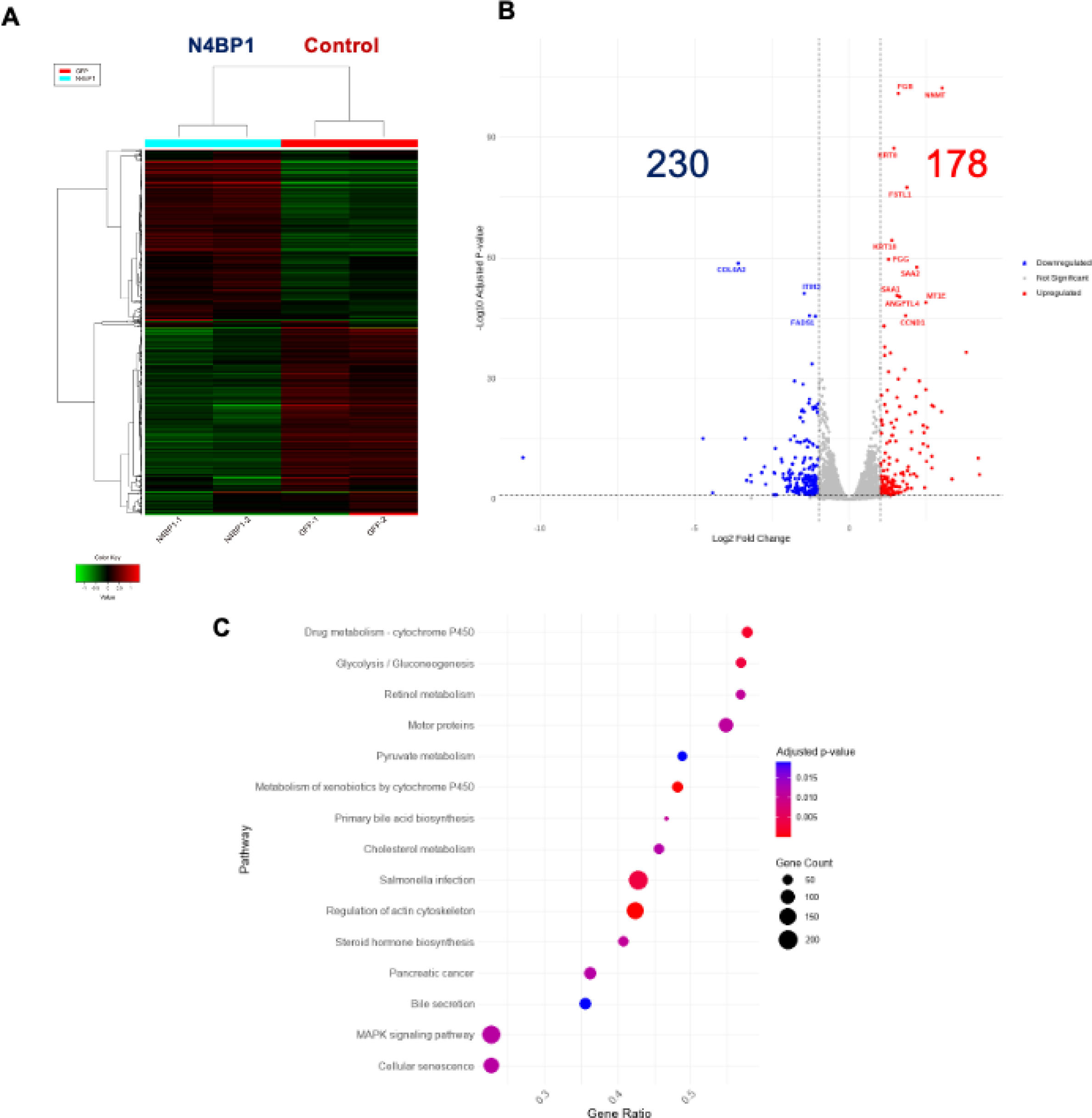
N4BP1 overexpression affected expression of several host factors. (A) Heat map of the transcriptomic profiles of N4BP1-expressing and control PHHs infected with HBV. (B) In the volcano plot, we set a cutoff value of gene expression at absolute fold change greater than 2 and an adjusted P-value less than 0.1 compared to control cells. Relative to the control cells, 178 factors were upregulated, and 230 factors downregulated in the N4BP1-overexpressing PHHs. (C) Pathway enrichment analysis of the differentially expressed genes in N4BP1-overexpressing PHHs versus the control PHHs demonstrates the distinct differences in pathways induced by N4BP1-overexpression. The graph depicts the number of genes corresponding to each pathway through the size of its dots. The color gradient, which transitions from red to purple, illustrates the adjusted p-value indicating significance of enrichment. The Gene Ratio on the horizontal axis indicates the proportion of input genes that are annotated to each term, thereby highlighting the distinct impact of N4BP1 over-expression. The raw data sets are shown in Supporting Information S1 table.

### N4BP1 expression is not induced by HBV

We then evaluated whether N4BP1 expression level was increased by HBV infection. A plasmid containing the 1.3-fold-overlength genome of HBV genotype C was transfected into Huh7 cells, and N4BP1 expression assessed by WB and qPCR at 3 d.p.t. Transfection of the HBV genome did not significantly affect N4BP1 expression at either the transcriptional or protein level (Fig 5A, 4B). N4BP1 expression also did not significantly change in HepG2-hNTCP C4 cells at 3 and 10 d.p.i. (Fig 5A, 5C). Finally, we measured N4BP1 expression in HepAD38.7 cells that stably secrete HBV particles unless cultured in the presence of tetracycline. In this system, removal of tetracycline and induction of HBV particle production also did not result in any change in N4BP1 expression (Fig 5A, 5D). Together, these results indicate that N4BP1 expression is not affected by HBV infection.

**Fig 5.**
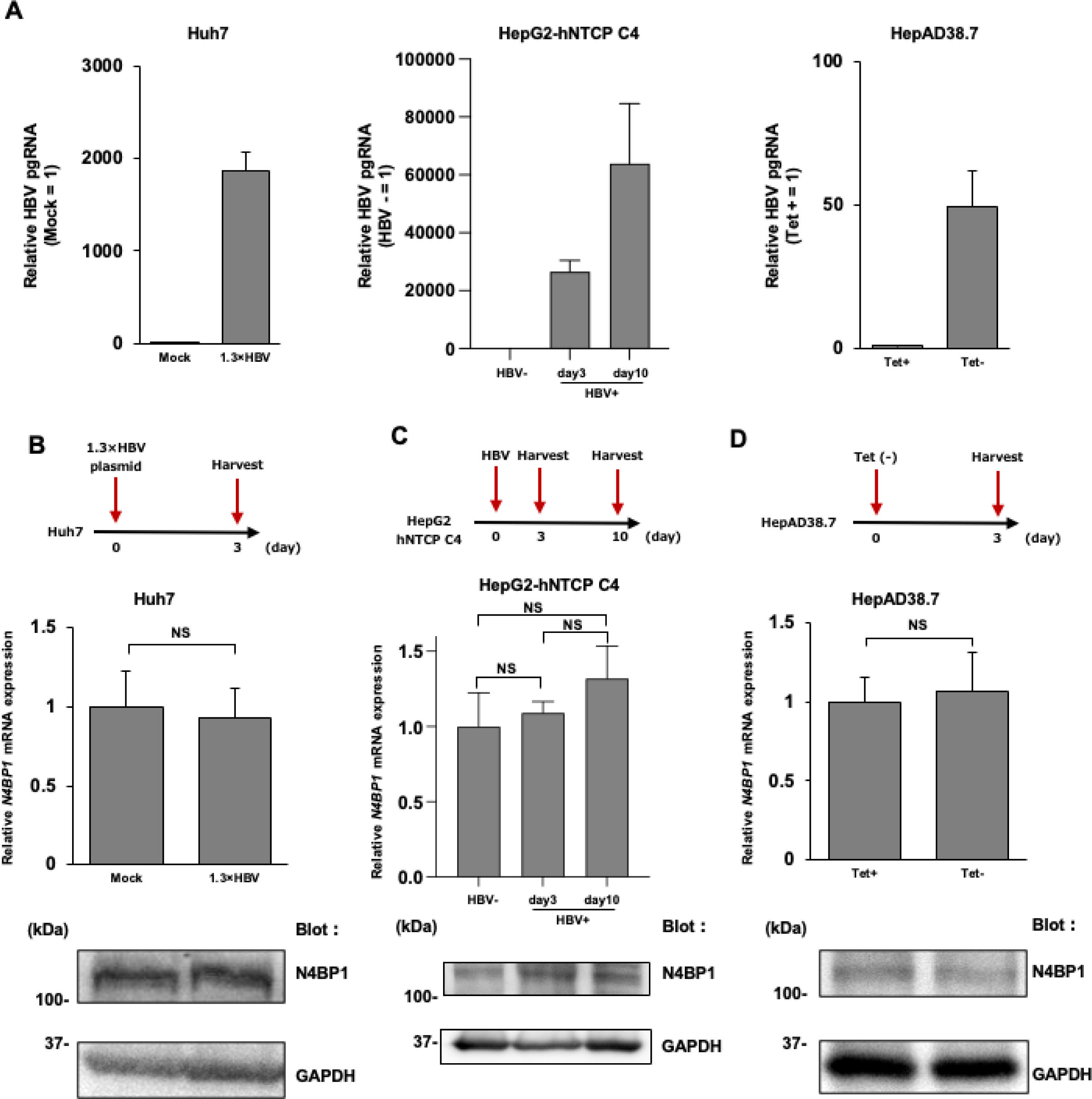
N4BP1 expression is not induced by HBV. (A) We evaluated whether N4BP1 expression is induced by HBV infection or transfection in hepatocyte cell lines. In each system, we confirmed by qPCR whether pgRNA was induced. (B) Huh7 cells were transfected with a plasmid containing the 1.3-fold-overlength genome of HBV genotype C and cells were collected at 3 d.p.t. (upper). *N4BP1* mRNA expression was assessed by qPCR (middle) and protein expression by WB (lower). (C) HepG2-hNTCP C4 cells were infected with HBV genotype D and cells were collected at 3 d.p.i. and 10 d.p.i. (upper). N4BP1 expression level by qPCR (middle). N4BP1 expression levels by WB (lower). (D) Tetracycline was removed from the medium of HepAD38.7 cells and the cells were collected 3 days later (upper). N4BP1 expression levels by qPCR (middle). N4BP1 expression levels by WB (lower). Error bars represent the SD. *P < 0.05, **P < 0.01, as determined by Mann-Whitney U test. All qPCR experiments were independently performed at least three times.

### The KH-like and RNase domains of N4BP1 are critical for the anti-HBV effect of N4BP1

As shown in Fig 6A, N4BP1 is composed of 896 amino acids and contains a KH-like domain at its N terminus side and an RNase domain at its C terminus (19). To examine which N4BP1 domains are needed to suppress HBV replication, we constructed deletion mutants lacking either the KH-like (ΔKH; deletion of amino acids 59-143) or the RNase domain (ΔRNase; deletion of amino acids 617-769).

**Fig 6.**
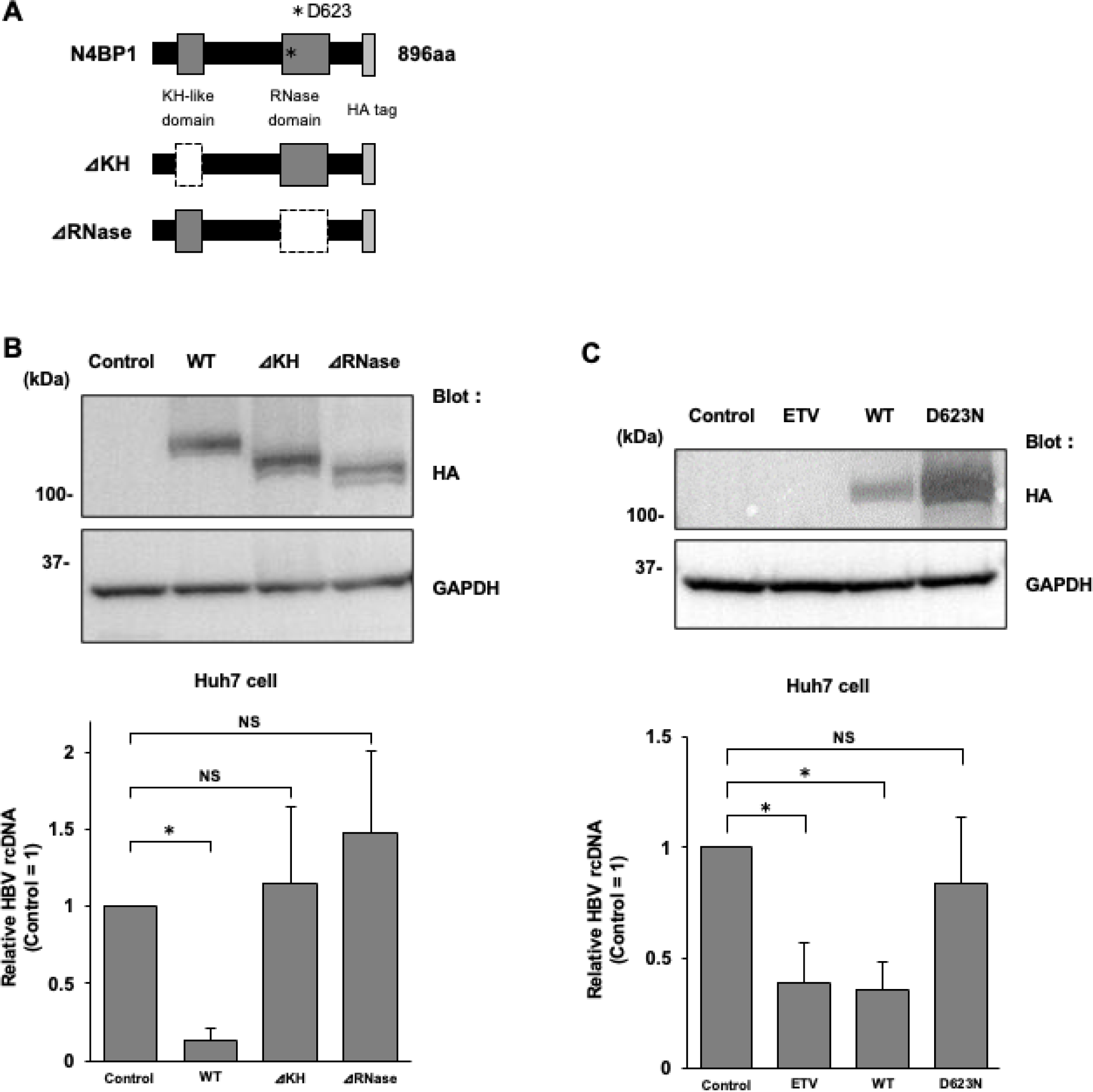
KH-like domain and RNase domain are involved in the anti-HBV effect of N4BP1. (A) Schematic of N4BP1. N4BP1 is composed of 896 amino acids and has both a KH-like domain and an RNase domain. We constructed a KH-like domain deficient mutant (deletion of amino acids 59-143) and an RNase domain deficient mutant (deletion of amino acids 617-769). Each plasmid had a HA tag at its C-terminus. (B) Huh7 cells were transfected with the plasmid containing the 1.3-fold-overlength genome of HBV genotype C and each expression plasmid (control vector, N4BP1 WT, ΔKH, and ΔRNase) and cells were collected at 3 d.p.t. WB was used to confirm the expression of each N4BP1 mutant (upper, anti-HA tag). GAPDH was used as a loading control (lower, anti-GAPDH). HBV levels were assessed by measuring intracellular rcDNA levels at 3 d.p.t. (C) Huh7 cells were transfected with the plasmid containing the 1.3-fold-overlength genome of HBV genotype C and each expression plasmid (control vector, WT, and D623N) and cells were collected at 3 d.p.t. WB was used to confirm the expression of each N4BP1 mutant (upper, anti-HA tag). GAPDH was used as a loading control (lower, anti-GAPDH). HBV levels were assessed by measuring intracellular rcDNA levels at 3 d.p.t. ETV was used as a positive control. Error bars represent the SD. *P < 0.05 as determined by Mann-Whitney U test. All qPCR experiments were independently performed at least three times.

The plasmid encoding wild-type or mutant N4BP1 was respectively co-transfected into naïve Huh7 cells with the plasmid containing the 1.3-fold-overlength genome of HBV genotype C and the rcDNA levels were measured. The expression of the deletion mutants was confirmed by WB using an HA-tagged antibody (Fig 6B, upper). The rcDNA level in N4BP1 overexpressing cells was significantly reduced, while the rcDNA levels in the ΔKH- or ΔRNase-expressing cells were comparable to those in the control cells (Fig 6B, lower).

Residue D623 in the RNase domain of N4BP1 (Fig 6A) has been shown to be important for the protein’s RNase activity (20). A D623N substitution causes loss of RNase activity without changing the structure of N4BP1. Co-transfecting the Huh7 cells with a plasmid expressing this N4BP1 D623 mutant and the HBV overlength genome also did not lead to a decrease in rcDNA levels (Fig 6C). These results suggest that both the KH-like and RNase domains are critical for the anti-HBV effect of N4BP1.

### N4BP1 promotes pgRNA degradation

Having shown the effect of N4BP1 on rcDNA, we then sought to determine more specifically which step of HBV replication N4BP1 suppresses. To that end, we assessed the impact of N4BP1 overexpression on cccDNA and pgRNA levels. HepG2-hNTCP C4 cells were infected with HBV and cells were harvested at 14 d.p.i. While cccDNA levels were not affected by overexpression of N4BP1, pgRNA and rcDNA levels were both significantly reduced (Fig 7A-7C). The HBV progeny secreted from infected HepG2-NTCP C4 cells is not infectious, and thus only single-round infection is observed in this system (21, 22). Thus, these results indicate that N4BP1 affects the steps of HBV replication after cccDNA was formed and augmented pgRNA degradation.

**Fig 7.**
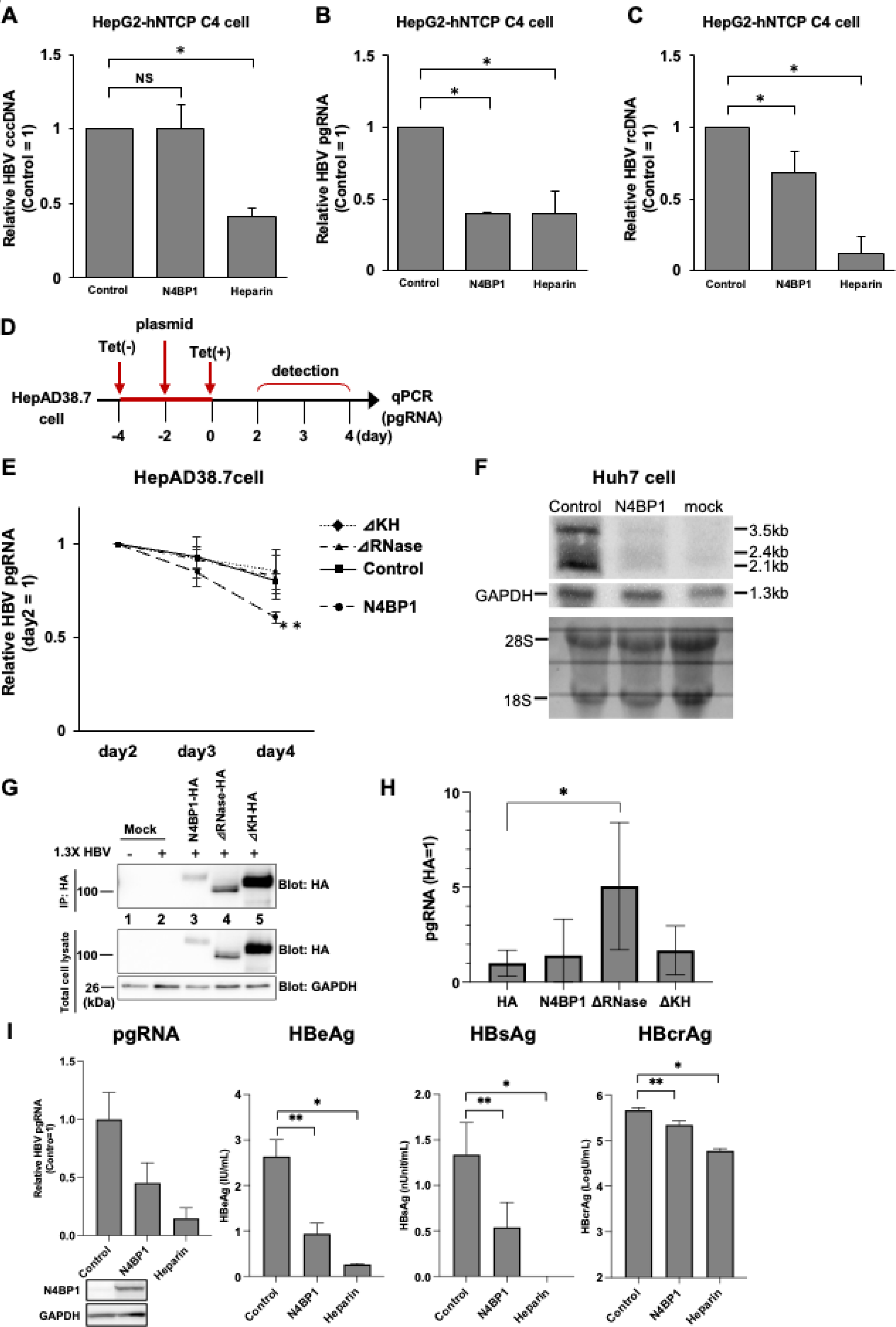
N4BP1 degrades HBV RNAs by binding HBV RNAs. HepG2-hNTCP C4 cells were infected with HBV genotype D and cells were collected at 14 d.p.i. Comparison of cccDNA (A), pgRNA (B), and rcDNA (C) levels in N4BP1-overexpressing HepG2-hNTCP C4 cells and control at 14 d.p.i. Heparin was used as a positive control. (D) Time course of pgRNA degradation assay in HepAD38.7 cells. (E) pgRNA levels over time in HepAD38.7 cells overexpressing N4BP1 WT, ΔKH, ΔRNase, or control. (F) Northern blotting analyses in control, N4BP1-overexpressing Huh7 cells transfected with the 1.3-fold-overlength genome of HBV and naïve Huh7 cells (mock). 28S and 18S ribosomal RNA was used as a loading control (lower panel). (G) Huh7 cells were co-transfected with the 1.3-fold-overlength genome of HBV genotype C with HA control vector, HA-tagged N4BP1 or ΔRNase. Cells were collected at 3 d.p.t. and RIP assay was performed. WB was used to confirm the expression of each N4BP1 mutant (Blot: HA) and IP was performed. GAPDH was used as a loading control (lower, anti-GAPDH). (H) RNA was extracted from total cell lysate and the immunoprecipitated protein, and then pgRNA was measured by qPCR. (I) PHHs were infected with HBV at 5 Geq/cell and supernatant was collected at 14 d.p.i. RNA was extracted from the cells and pgRNA was measured and N4BP1 overexpression was confirmed by western blot (left). The viral protein levels in supernatant were measured by HBeAg CLIA (second from left), HBsAg ELISA (second from right) and HBcrAg assay (right). Error bars represent the SD. *P < 0.05, **P < 0.01, as determined by Mann-Whitney U test. All experiments were independently performed at least three times.

To elucidate the mechanism of pgRNA suppression by N4BP1, we turned to HepAD38.7 cells. First, the cells were cultured for 2 days in media without tetracycline to induce HBV replication. Then N4BP1 or the domain deletion mutant plasmids were transfected, and tetracycline added at 2 d.p.t. to stop viral replication. Cells were collected starting 2 days after adding tetracycline to measure the pgRNA levels (Fig 7D). As shown in Fig 7E, the expression of wild-type N4BP1 augmented the rate of pgRNA degradation. In contrast, both domain deficient mutants resulted in pgRNA degradation at a rate similar to the control, suggesting that N4BP1 affects the degradation rate of pgRNA. Next, we examined whether N4BP1 expression also suppresses HBV RNAs other than pgRNA. We co-transfected Huh7 cells with a plasmid expressing N4BP1 and a plasmid containing 1.3-fold-overlength genomes of HBV genotype C. At 3 d.p.t., cells were collected and HBV RNA expression was measured by Northern blotting. Results showed that control Huh7 cells showed intense expression of 3.5kb, 2.4/2.1kb HBV RNA, whereas N4BP1 overexpression Huh7 cells showed weak expression of all these RNAs (Fig 7F).

Next, to test that N4BP1 binds HBV RNA directly, we performed an RNA immunoprecipitation (RIP) assay. Huh7 cells were co-transfected with a plasmid containing the 1.3-fold-overlength genome of HBV genotype C and HA-tagged N4BP1 wildtype or domain deficient mutants or HA control vector. Cells were collected at 3 d.p.t. and immunoprecipitated with anti-HA-tag monoclonal antibody magnetic beads. Wildtype and N4BP1 domain-deficient mutants were confirmed to be precipitated by an anti-HA-tag monoclonal antibody by western blot (Fig. 7G). The RNA-protein interaction was expected to be most stable in the ΔRNase domain mutant, and our RIP-quantitative PCR (qPCR) assay confirmed this (Fig. 7H). In contrast, the amount of pgRNA recovered with wildtype N4BP1 and ΔKH-like domain mutants was far less, presumably due to the intact RNase activity.

Furthermore, to investigate whether N4BP1 also led to reduced expression of viral proteins, we assessed levels of HBeAg, HBsAg and HB core-related antigen (HBcrAg) in the supernatant of HBV-infected PHHs at 14d.p.i., the timepoint when we had observed decreased pgRNA (Fig. 7I). All 3 HBV proteins we evaluated were decreased in the N4BP1-overexpressing PHHs compared to control, indicating that N4BP1 inhibits pcRNA, PreS2/S RNA and potentially pgRNA, consistent with our northern blot data.

### N4BP1 expression is induced by IFN-α/λ in PHHs

It has been reported that N4BP1 expression is increased 3- to 5-fold with IFN-α treatment in Jurkat cells and THP-1 cells (20). To see if this was the case in PHHs and hepatoma cell lines, we treated Huh7, HepG2-hNTCP C4, and PHHs with 1000 U/ml of IFN-α and 100 ng/ml of IFN-λ, which are known to also inhibit HBV replication. Cells were collected at 6, 24 and 48 hours post-drug treatment without HBV infection, and expression of *N4BP1* mRNA and protein was examined by qPCR and WB, respectively (Fig 8A). *ISG15* mRNA was included as a positive control. As shown in Fig 8B, levels of *N4BP1* mRNA were increased 2-fold in both Huh7 and HepG2-hNTCP C4 cells by IFN-α at 6 hours and 24 hours, respectively, which is small, albeit significant in expression. IFN-λ stimulation increased levels of *N4BP1* mRNA in PHHs at 24 hours and 48 hours significantly and N4BP1 protein levels were prompted in Huh7 cells and PHHs (Fig 8C).

**Fig 8.**
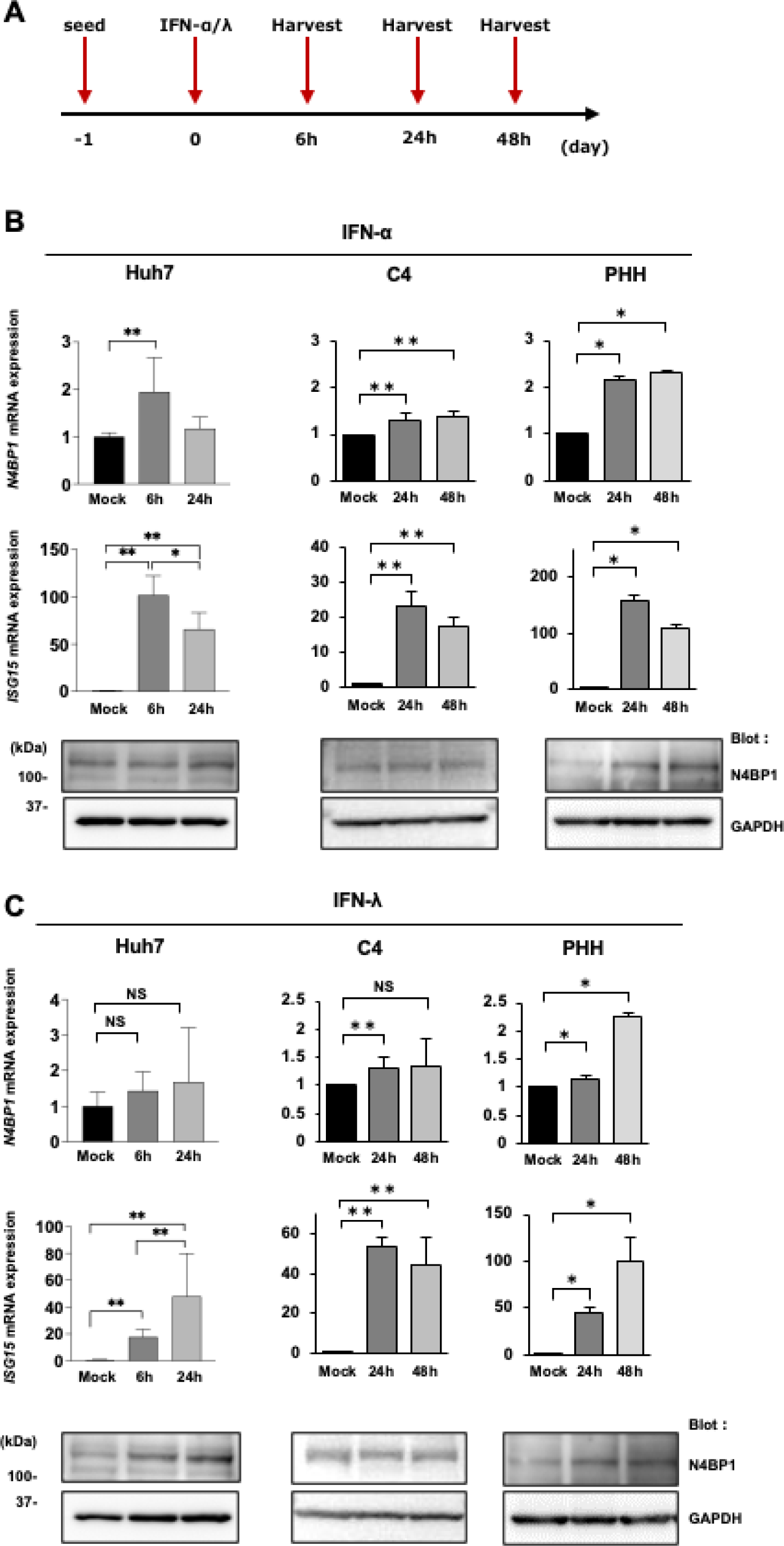
N4BP1 expression is induced by IFN-α and IFN-λ in PHHs. (A) Time course of the drug treatment assay. Cells were harvested at 6, 24 or 48 hours post-IFN-α/λ treatment. (B) N4BP1 expression levels by qPCR in Huh7 cells, HepG2-hNTCP C4 cells, and PHHs treated with IFN-α (upper). ISG15 expression levels by qPCR (middle). N4BP1 protein expression levels by WB. GAPDH was used as a loading control (lower). (C) N4BP1 expression levels by qPCR in Huh7 cells, HepG2-hNTCP C4 cells, and PHHs treated with IFN-λ (upper). ISG15 expression levels by qPCR (middle). N4BP1 protein expression levels by WB. GAPDH was measured as loading control (lower).

IFN-α and IFN-λ treatment in PHHs resulted in an approximately 2-fold increase in *N4BP1* mRNA expression and protein level was also increased. Collectively, these results indicate that IFN-α treatment induces N4BP1 expression in Huh7 and HepG2-hNTCP C4 cells, while both IFN-α and IFN-λ induces N4BP1 expression in PHHs. As primary cells, PHHs are more physiologically relevant than hepatoma lines such as Huh7 and HepG2-hNTCP C4 cells in which the response to IFN is disrupted. To build upon our in vitro findings, we then examined samples from patients with HBV-HCC to assess whether there was an association between *N4BP1* mRNA expression in the liver and HBV reactivation status. All the patients showed HBs-Ag (+), HBe-Ag (-) and HBe-Ab (+), and categorized as either HBV DNA positive or negative per HBV DNA levels in the serum, and *ISG15* and *N4BP1* mRNA expression were assayed in non-cancerous liver tissue from these patients (Fig. 9). *ISG15* mRNA did not significantly differ between the patient groups, but *N4BP1* mRNA was significantly higher in patients who were HBV DNA negative vs positive.

**Fig 9.**
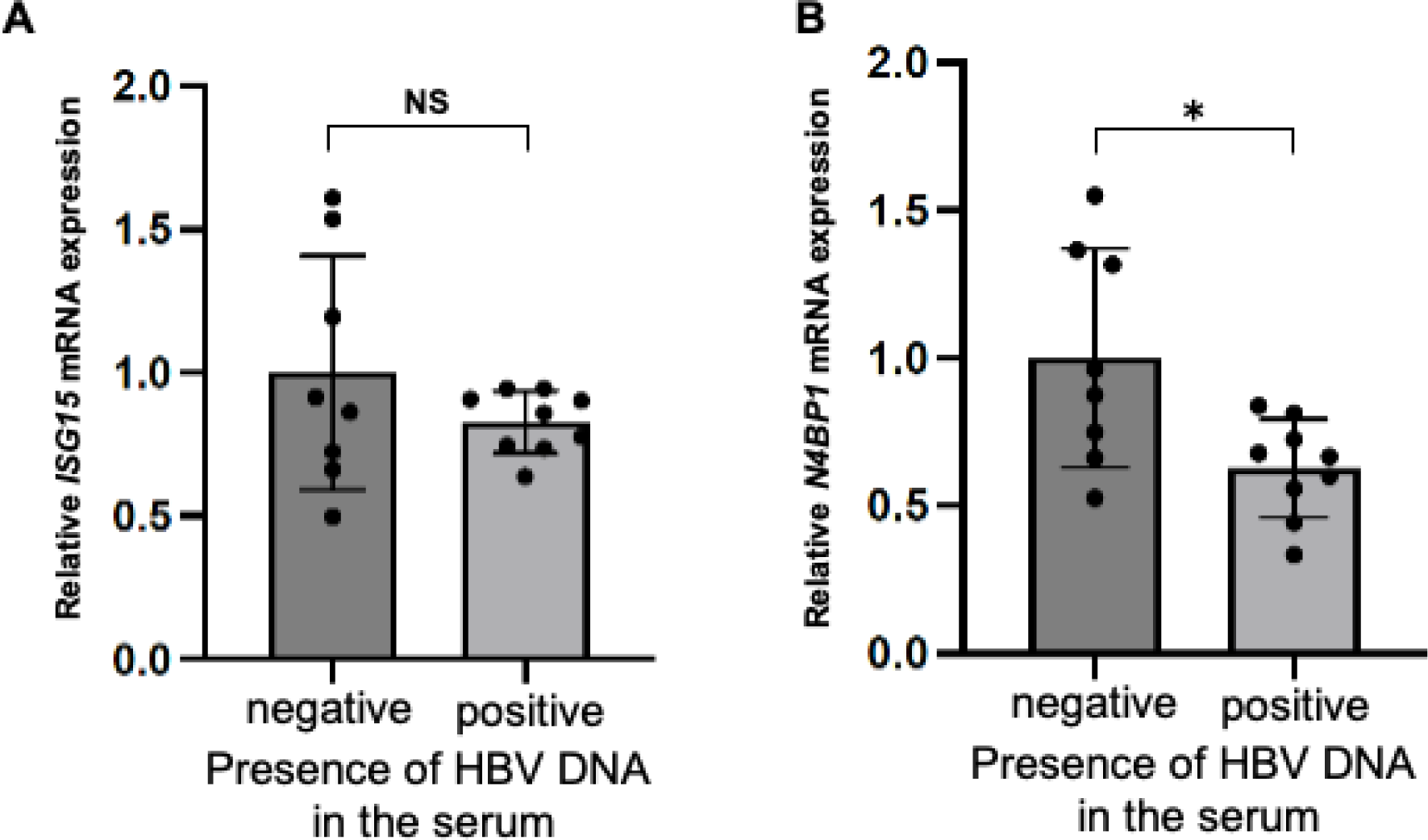
ISG15 and N4BP1 expression levels in HBV-infected human liver tissues. The mRNA levels of *ISG15* (A) and *N4BP1* (B) from non-cancerous tissues of HBV-HCC patients with HBs-Ag (+), HBe-Ag (-), HBe-Ab (+) were examined by qPCR. The samples were divided into HBV DNA positive group (n=9) and HBV DNA negative group (n=8). Error bars represent the SD. *P < 0.05, as determined by Mann-Whitney U test.

## Discussion

NAs are generally well tolerated, very effective in inhibiting HBV replication, and are currently the most commonly used treatment for chronic hepatitis B. However, since NAs inhibit reverse transcription, HBV cccDNA remains in the nucleus of hepatocytes so the patient is never completely cured. Indeed, patients who discontinue NA treatment can experience virologic relapse and thus must continue NA therapy for extended periods. Long-term treatment with NA is associated with the development of antiviral drug resistance; for example, mutations in the HBV polymerase gene, which is a target of NAs, have been reported (23-25). Resistance to NA therapy can lead to hepatitis relapse and even death. Thus, we sought to identify host factors involved in HBV proliferation that would be distinct from the viral sites targeted by NAs. In a library of 132 RBPs containing proteins with known RNA-binding domains such as RNA recognition motif domains or zinc finger domains, we found that N4BP1 inhibited HBV replication, was negatively correlated with HBV rcDNA levels in hepatoma cell lines and suppressed HBV replication by promoting pgRNA degradation.

We found that both the KH-like domain and the RNase domain of N4BP1 were essential for the protein’s anti-HBV effect (Fig 6B). It has been reported that the KH-like domain binds to RNA or single-strand DNA and is involved in splicing, transcriptional regulation, and translational regulation (19, 26, 27). On the other hand, the RNase domain of N4BP1 is also involved in regulating genome stability and repressing the expression of target genes through mRNA strand breaks (28). Based on our RIP assay data, it is possible that N4BP1 binds HBV RNA via its KH-like domain and degrades HBV RNA by the RNase activity of its RNase domain. However, this would need to be confirmed with a RIP assay using a doubly deficient RNase and KH-like domain mutant. In fact, in this study, N4BP1 suppressed HBV replication at the step of pgRNA production in the HBV life cycle (Fig 7). We also found that N4BP1 suppresses production of not only the 3.5 kb RNA of HBV, but also the 2.4/2.1 kb RNA by northern blot, which implies N4BP1 promotes degradation of pgRNA, PreS1 and PreS2/S. However, it is still unclear and remains to be assessed whether HBx is also affected by N4BP1.

N4BP1 is known to be an inhibitor of innate immunity and inflammation-mediated cytokine production. NF-κB is an important transcription factor for the elicitation of cytokine responses, and its activation is necessary for many immune responses. N4BP1 interacts with the NF-κB signal essential modulator (NEMO, also known as IκB kinase γ) to suppress Toll-like receptor (TLR)– dependent NF-κB activation (29). Furthermore, even in vivo, it has been reported that N4BP1-/- mice develop mild inflammation and have exacerbated TLR-dependent inflammatory responses (30). Recently, N4BP1 was found as a new hotspot gene for HBV integration (31). Although the significance of HBV integration to N4BP1 remains unknown, it is interesting that the virus integration occurred to N4BP1 which has anti-HBV activity. Regarding virus replication, N4BP1 is known to directly recognize and degrade retroviruses such as HIV-1 (20) and to be involved in replication of the Porcine Reproductive and Respiratory Syndrome Virus (32). In both cases, the RNase activity of N4BP1 was found to inhibit viral replication by directly degrading viral RNA, consistent with our data. In addition, the RNase domain of N4BP1 has a similar structure to that of MCPIP1(33), which has been reported to recognize and degrade HBV pgRNA directly (15). However, MCPIP1 is a cytoplasmic RNase, whereas N4BP1 is mainly localized on the nucleus. Furthermore, MCPIP1 has not been reported to suppress 2.4 kb and 2.1 kb HBV RNAs, whereas we have demonstrated that N4BP1 suppresses these RNAs. These results suggest that MCPIP1 and N4BP1 may regulate HBV degrading mRNAs through different processes. The mechanism behind our finding that the KH-like domain of N4BP1 was also important for the anti-HBV effect of this protein is presumably due to its binding to HBV RNA but further work is needed to elucidate the mechanism more precisely.

Although N4BP1 expression has been reported to be induced by type I IFN (20), stimulation of N4BP1 expression by IFN-α (type I IFN) and IFN-λ (type III IFN) treatment varied between the PHHs and hepatoma cell lines we tested. This could be due to the disrupted IFN signaling reported in the cell lines Huh7 and HepG2 that result in them being less sensitive to IFN (34, 35). Also, ISG15 is an inducible factor by IFN-α and IFN-λ (36, 37). In characterizing N4BP1 expression, we found that N4BP1 expression was not induced by HBV infection or transfection in hepatoma cell lines (Fig 5). Furthermore, in clinical liver tissue samples, N4BP1 expression was significantly lower in HBV-DNA positive vs HBV-DNA negative samples (Fig 9B). These results suggest that N4BP1 expression may differ according to each individual and that expression of N4BP1 in the liver may impact HBV reactivation. Otherwise, it is thought that factors upstream of N4BP1 may cause differences in N4BP1 expression and affect HBV reactivation. In addition, research has shown that N4BP1 can be inactivated by caspase-8 (30) or mucosa-associated lymphoid tissue lymphoma translocation 1 (MALT1) (20) in immune cells. In the latter instance, N4BP1 inactivation by MALT-1 induced HIV-1 reactivation. This does not appear to be the case in our study, as our RNA-seq data did not indicate MALT-1 induction by N4BP1 overexpression in PHH, but other signaling proteins may have a similar effect, resulting in greater HBV replication in hepatocytes. This will need to be studied further.

A limitation of our current study is the small number of patient samples used to evaluate the relationship between N4BP1 and HBV reactivation. Studies with a greater number of samples are needed to further characterize this possible relationship. In addition, although transcriptomic analysis implies that N4BP1 was not associated with transcriptional regulation, it is required to strengthen this result by other methods such as ActD chase assay. Also, identification of the element(s) of HBV RNA that bind N4BP1 remains to be determined. As HBV RNAs overlap each other, explicit identification of the binding region(s) was difficult. A previous report suggested that sequence motif(s) other than GC dinucleotides may be recognized by N4BP1 as N4BP1 can degrade HIV-1 transcripts, which have a markedly reduced frequency of GC dinucleotides (20). However, based on our data, it is conceivable that an overlapping area or areas of the 3.5kb and 2.4kb/2.1kb HBV RNAs may bind to N4BP1.

In conclusion, to investigate different treatments against HBV infection from currently approved HBV therapeutics, we focused on RBPs. We found that N4BP1 is a RBP with novel anti-HBV activity, inhibiting HBV replication via degradation of HBV RNAs. Although the detailed molecular mechanism behind the anti-HBV effect of N4BP1 remains to be uncovered, understanding the association between N4BP1 and HBV could contribute to the development of new HBV treatments.

## Materials and methods

### Cell culture

All cells were grown at 37℃ in a 5% (v/v) CO2 environment. Huh7 cells and HEK293T cells were obtained from the Japanese Collection of Research Bioresources Cell Bank (JCRB, Osaka, Japan, JCRB0403 and JCRB9068). HepG2-hNTCP C4 cells and HepAD38.7 cells were kindly provided by Takaji Wakita (NIID, Tokyo, Japan). PHHs were obtained from Phoenix Bio (Hiroshima, Japan). Huh7 cells and HEK293T cells were maintained in DMEM (Nacalai Tesque, Kyoto, Japan) supplemented with 100 U/ml of penicillin, 100 μg/ml of streptomycin (Nacalai Tesque) and 10% fetal bovine serum (FBS, Sigma, MO, USA). HepG2-hNTCP C4 cells were maintained in the aforementioned media containing 400 μg/ml G418 (Nacalai Tesque). HepAD38.7 cells were maintained in DMEM/Ham’s F-12 media (Nacalai Tesque) supplemented with 10% FBS, 100 U/ml of penicillin, 100 μg/ml of streptomycin, 400 μg/ml of G418, 5 μg/ml of insulin (Nacalai Tesque), and 400 ng/ml of tetracycline (Nacalai Tesque). PHHs were maintained in PHH-specific media provided by the manufacture (Phoenix Bio).

### Generation of RBP overexpressing cells

RBP-overexpressing Huh7 cells were generated by transfection of RBP expression plasmids using TransIT LT-1 transfection reagent (Mirus, WI, USA). Briefly, Huh7 cells were seeded in 24-well plates (Greiner, Bad Haller, Austria) 1 day prior to the transfection. The following day, 1 μl of TransIT LT-1 transfection reagent, 0.3 μg of RBP expression plasmid, and 0.1 μg of a plasmid containing a 1.3-fold-overlength genome of HBV (genotype C) were mixed in 50 μl of Opti-MEM (Thermo Fisher Scientific, MA, USA). After 15 minutes incubation at room temperature, the mixture was added into the supernatant of the cultured Huh7 cells. Entecavir (Fujifilm, Tokyo, Japan) was added at 5 μM the day after transfection as a positive control.

RBP-overexpressing HepG2-hNTCP C4 cells were made by lentiviral transduction. Lentiviruses were made by transfecting HIV gag/pol, VSV-G, REV, and pcs-II EF RBP expression plasmids into HEK293T cells using TransIT LT-1 transfection reagent. At 2 d.p.t., the supernatant of HEK293T was collected and spun at 4,200 rpm for 2 minutes at room temperature and the supernatant added to HepG2-hNTCP C4 cells. Entecavir or heparin was used as a positive control. Entecavir was added at 5 μM the day after infection. Heparin (Mochida Pharmaceutical Co., Tokyo, Japan) was added at 5 U/ml 30 min before infection.

To express N4BP1 in PHHs, we used an adenovirus vector (AdV). The N4BP1expressing adenovirus vector (N4BP1 AdV) was constructed using the cosmid cassette pAxEFwit2, which has a cloning site *SwaI* flanked by the EF1α promoter and rabbit β-globin polyA signal in the E1-deleted region of the adenovirus genome. The N4BP1 cDNA was prepared from the pCS2-EF-N4BP1-HA plasmid and inserted into the *SwaI* cloning site. The AdV was prepared as previously described (38, 39). The AdV was titrated using the methods described by Pei et al (40). Briefly, the copy numbers of viral genome successfully transduced into target cells were measured by qPCR. For generating N4BP1-overexpressing PHHs, cells were infected with N4BP1 AdV at a multiplicity of infection (MOI) of 5 or 30 two days before HBV infection.

### Generation of RBP knockdown cells

RBP knockdown cell lines were generated using Silencer Select siRNA (Thermo Fisher Scientific) and Lipofectamine RNAiMAX (Thermo Fisher Scientific) following the manufacturer’s protocol. Briefly, 1.5 μl of Lipofectamine RNAiMAX and 5 pg of siRNA were mixed in 50 μl of Opti-MEM. After incubating for 5 minutes at room temperature, the mixture was gently added to the cell supernatant. The following siRNA reagents were used in this study: control siRNA (Silencer Select Negative Control No.1 siRNA, 4390843), ZCCHC10 siRNA#1 (s29507), ZCCHC10 siRNA#2 (s29508), ZCCHC10 siRNA#3 (s29509), HNRNPC siRNA#1 (s6719), HNRNPC siRNA#2 (s6720), HNRNPC siRNA#3 (s6721), N4BP1 siRNA#1 (s18638), N4BP1 siRNA#2 (s18639).

### Generation of N4BP1 knockout cells

N4BP1 KO Huh7 cells were generated by CRISPR/Cas9 using pX330-U6-Chimeric_BB-CBh-hSpCas9 (pX330, Addgene, MA, USA) and HR110PA-1 (System Biosciences, CA, USA). The pX330-puro vector was constructed by integrating the puromycin resistance gene sequence from the HR110PA-1 vector into the pX330 vector. The pX330-puro vector was digested using *BbsI* (Biolabs, MA, USA) and the target sequence was subcloned. The plasmid was transfected into Huh7 cells using TransIT LT-1 transfection reagent and after selection with puromycin dihydrochloride (Invivogen, CA, USA), a single colony was isolated using cloning cylinders (Merck, Darmstadt, Germany) and propagated. KO of the target gene was confirmed by WB and DNA sequencing.

### Plasmid Constructions

We used the following plasmids containing 1.3-fold-overlength genomes of different HBV genotypes: pUC19-Ae_us (genotype A, accession no. AB246337), pGEM-Bj-JPN56 (genotype B, accession no. AB246342), and pUC19-C_JPNAT (genotype C, accession no. AB246345).

Each RBP expression plasmid had a C-terminus HA tag and was inserted between the *XhoI* and *XbaI* sites of the pcs-II EF vector. The KH-like domain deletion mutant (ΔKH), RNase domain deletion mutant (ΔRNase), and D623N mutant were generated from the N4BP1 expression plasmid. ΔKH plasmid was generated by deleting amino acids 59-143 from N4BP1; ΔRNase was generated by deleting amino acids 617-769 N4BP1; and D623N was generated by changing the 623rd amino acid from asparagine acid to asparagine. To insert these mutated N4BP1 genes into the pcs-II EF vector between the *XhoI* and *XbaI* sites, we used the following primer sets and In-Fusion Snap Assembly Master Mix (Takara Bio, Shiga, Japan) per the manufacturer’s directions. Each N4BP1 mutant plasmid had a C-terminus HA tag.

ΔKH: 5’- CGCTACCGGTCTCGAGATGGCGGCCCGGGCGGTGCT -3’ (forward) 5’- GGGTAGGTTCTCTTTCGCCCCGCAGAGCTGCAGCC -3’ (reverse), and 5’- CAGCTCTGCGGGGCGAAAGAGAACCTACCCAGTAG -3’ (forward) 5’- GGTACATGGTCTCGAGATCCAACACCATGGCAGAAA -3’ (reverse)

ΔRNase: 5’- CGCTACCGGTCTCGAGATGGCGGCCCGGGCGGTGCT -3’ (forward) 5’- TTCCTTCTGAAGAAAATCCGTTCTCCCTGGTTCAT -3’ (reverse), and 5’- CCAGGGAGAACGGATTTTCTTCAGAAGGAAGTCTG -3’ (forward) 5’- GGGTAGGTTCTCTTTCGCCCCGCAGAGCTGCAGCC -3’ (reverse)

D623N: 5’- CGCTACCGGTCTCGAGATGGCGGCCCGGGCGGTGCT -3’ (forward) 5’- AACATTGCTCCCATTTATAACAATGTGTTTCAAAT -3’ (reverse), and 5’- AATGGGAGCAATGTTGCAATTACCCATGGTCTGAA -3’ (forward) 5’- GGTACATGGTCTCGAGATCCAACACCATGGCAGAAA -3’ (reverse)

### Viruses

The production of HBV particles (genotype D) was adapted from a previous report (41). Briefly, HepAD38.7 cells were cultured under tetracycline (400 ng/mL) for maintenance. To produce HBV particles, tetracycline was removed and the supernatant collected after 6 and 9 days. The collected supernatant was mixed with 30% polyethylene glycol (PEG) 8000 (Sigma) and incubated overnight at 4℃. Then it was spun at 3,000 rpm for 20 minutes at 4℃ for concentration. Before HBV infection, HepG2-hNTCP C4 cells were seeded in collagen type I-coated 24-well plates (Iwaki, Tokyo, Japan) and transduced with lentivirus to express RBP (see “Generation of RBP-overexpressing cells” above). At 2 days post-transduction, the cells were infected with 5,000 genome equivalents (GEq)/cell of HBV in DMEM containing 3% DMSO (Nacalai Tesque) and 4% PEG 8000, and the media was changed every 2-3 days until 14 d.p.i..

HBV genotype C for PHHs infection was purchased from Phoenix Bio. The infection into PHHs was performed at 5 Geq/cell in PHH-specific media containing 40% PEG 8000 following the manufacturer’s protocol. The culture media was replaced every 4 days. Cells were harvested at 14 d.p.i. for WB and reverse transcription quantitative PCR (RT-qPCR).

### Western blotting

Cells were lysed with 2 × Laemmli sample buffer (Bio Rad Laboratories, CA, USA) containing 10% 2-mercaptoethanol (Sigma) and boiled for 10 minutes at 100℃. Then lysates were loaded on a 10-20% gradient SDS-PAGE gel (Atto, Tokyo, Japan) and transferred onto a polyvinylidene fluoride transfer membrane (Millipore, MA, USA). Membranes were immersed in Block Ace (Kac, Kyoto, Japan) containing 1% bovine serum albumin (Sigma) and incubated for 1 hour at room temperature. The blocking solution was then decanted, and replaced with Can Get Signal Solution 1 (Toyobo, Osaka, Japan) containing the appropriate antibodies described below and incubated for 1 hour. The membranes were washed with 1 × PBS (Fujifilm) containing 0.1% polyoxyethylene (20) sorbitan monolaurate (Fujifilm) and incubated with Can Get Signal Solution 2 (Toyobo) containing horseradish peroxidase-conjugated antibody against rabbit or mouse immunoglobulins for 1 hour. After addition of Amersham ECL Prime Western Blotting Detection Reagents (Cytiva, Tokyo, Japan), the membranes were subsequently imaged via LuminoGraph I (Atto).

The following antibodies were used in this study: anti-N4BP1 (Bethyl Laboratories, TX, USA), anti-ZC3H12B (Gene Tex, CA, USA), anti-HNRNPC (abcam, Cambridgeshire, UK), anti-ZCCHC10 (Sigma), anti-KHNYN (Fujifilm), anti-HA tag (Biolegend, CA, USA), anti-rabbit IgG (H+L) HRP (Jackson Immuno Research Laboratories, MD, USA), anti-mouse IgG (H+L) HRP (Thermo Fisher Scientific), and anti-GAPDH (Fujifilm). The antibodies used in this study are listed in S2 Table.

### Purification of intracellular HBV rcDNA

To extract intracellular HBV DNA, cells were washed twice with PBS, and cells were lysed in lysis buffer (100 mM Tris-HCl [pH 8.0], 0.2% NP-40) for 15 minutes at 4℃. After spinning at 13,000 rpm for 1 minute at room temperature, the supernatant was collected and incubated with 1 M MgCl2, 0.2 mg/ml of DNase I (Sigma), and 10 mg/ml of RNase A (Nacalai Tesque) for at least 3 hours at 37℃. After adding 500 mM EDTA and 5 M NaCl, the lysates were digested with 20 mg/ml of proteinase K (Kanto Chemical, Tokyo, Japan) and 10% SDS for 5 hours at 55℃. The extracted HBV DNA was purified using phenol-chloroform-isoamyl alcohol (Sigma), then precipitated with isopropanol, 0.5 mg/ml of glycogen, and 3M sodium acetate. After washing with 70% ethanol, the purified HBV DNA was resolved in 50 μl pure water. HBV rcDNA was detected by qPCR using Power SYBR Green PCR Master Mix (Applied Biosystems, CA, USA). As a standard for rcDNA, pUC19-C_JPNAT was used. qPCR procedures were following the manufacturer’s protocol. Briefly, the samples were incubated at 50℃ for 2 minutes and at 95℃ for 10 minutes to initial denaturation. Then, the samples were incubated at 95℃ for 15 seconds and at 60℃ for 1 minute for 40 cycles to polymerase reaction. The following primer pair was used in this study.

rcDNA: 5’- GGAGGGATACATAGAGGTTCCTTGA -3’ (forward) 5’- GTTGCCCGTTTGTCCTCTAATTC -3’ (reverse)

### Quantification of intracellular HBV cccDNA level

To extract intracellular HBV cccDNA, the cells were lysed by lysis buffer (10mM Tris-HCl [pH 7.4], 10mM EDTA, 0.5% SDS) and 5M NaCl was added. After incubation overnight at 4℃, the samples were spun at 15,000 rpm for 30 minutes at 4℃ and the supernatant was collected to isolate protein-free DNA. The DNA was purified a total of three times by phenol (Nacalai Tesque), phenol-chloroform-isoamyl, and chloroform (Fujifilm). After precipitation with isopropanol and ethanol, Plasmid-Safe ATP-Dependent DNase (Lucigen, WI, USA) was added to the purified DNA to remove double-stranded and circular DNA. The Purified DNA was extracted in 30 μl of 10mM Tris-Cl (pH 8.5) using the FastGene Gel/PCR Extraction Kit (NIPPON Genetics, Tokyo, Japan). HBV cccDNA was measured by qPCR using TaqMan Fast Advanced Master Mix (Applied Biosystems, CA, USA). As a standard for cccDNA, pUC19-C_JPNAT was used. qPCR procedures were done per the manufacturer’s protocol. In brief, the samples were incubated at 50℃ for 2 minutes and at 95℃ for 10 minutes to initiate denaturation. Then, the samples were incubated at 95℃ for 15 seconds and at 60℃ for 1 minute for 50 cycles for the polymerase reaction. The following primer pair and probe were used in this study.

cccDNA: 5’- CGTCTGTGCCTTCTCATCTGC -3’ (forward) 5’- GCACAGCTTGGAGGCTTGAA -3’ (reverse) 5’- CTGTAGGCATAAATTGGT -3’ (Probe)

### Measurement of intracellular HBV pgRNA

Total RNA including pgRNA was extracted using the RNeasy Mini Kit (Qiagen, Hilden, Germany) following the manufacturer’s protocol. The extracted RNA was resolved in 50 μl pure water and measured by RT-qPCR using Power SYBR Green RNA-to-Ct 1-step Kit (Applied Biosystems). The pgRNA expression level was normalized against the expression level of GAPDH as a housekeeping gene and calculated by the ΔΔCT method. RT-qPCR procedures were done following manufacturer’s protocol. First, the samples were incubated at 50℃ for 2 minutes and at 95℃ for 10 minutes to intiate denaturation. Then, the samples were incubated at 95℃ for 15 seconds and at 60℃ for 1 minute for 40 cycles for the polymerase reaction. Samples were then incubated at 95℃ for 15 seconds, 60℃ for 1 minute, and 95℃ for 15 seconds for the melting curve measurement. The following primer pairs were used in this study.

pgRNA: 5’- TCCCTCGCCTCGCAGACG -3’ (forward) 5’- GTTTCCCACCTTATGAGTC -3’ (reverse)

GAPDH: 5’- AGAAGGCTGGGGCTCATTTG -3’ (forward) 5’- AGGGGCCATCCACAGTCTTC -3’ (reverse)

### pgRNA degradation assay

HepAD38.7 cells were seeded in collagen type I-coated 12-well plates (Iwaki) and cultured overnight in media containing tetracycline hydrochloride. After washing twice with PBS, cells were cultured without tetracycline for 48 hours. Then, the cells were transfected with N4BP1 wild-type expression plasmid, ΔKH expression plasmid, ΔRNase expression plasmid, or control vector. These plasmids were transfected as follows: 100 μl of Opti-MEM, 6 μl of TransIT LT-1, and 3μg of expression plasmid. After transfection of plasmids, cells were cultured for 48 hours and changed to a tetracycline-containing media in order to stop HBV production. Cells were collected at 2, 3, and 4 days after changing to tetracycline-containing media. Total RNA, including pgRNA, was extracted as described above for RT-qPCR.

### Drug treatment assay

Huh7 cells, HepG2-hNTCP C4 cells, and PHHs were stimulated with 1000 U/ml of IFN-α (PBL Assay Science, NJ, USA) or 100 ng/ml of IFN-λ (Pepro Tech, NJ, USA). At 6, 24 and 48 hours post-drug stimulation, total RNA in cells was extracted for qPCR using the RNeasy Mini Kit, and cDNA was synthesized using SuperScript IV VILO Master Mix (Invitrogen). For WB, the cells were lysed with 2 × Laemmli sample buffer containing 10% 2-mercaptoethanol and processed as described above. qPCR was performed using Power SYBR Green PCR Master Mix. The mRNA expression levels of N4BP1 and ISG15 were normalized against the expression level of GAPDH as a housekeeping gene and calculated by the ΔΔCT method. The following primer pairs were used in this study.

N4BP1: 5’- CCCGATGATCCTCTGGGAAG -3’ (forward) 5’- TTTGGCAGGGCACTGAGTAG -3’ (reverse)

ISG15: 5’- ACTCATCTTTGCCAGTACAGGAG -3’ (forward) 5’- CAGCATCTTCACCGTCAGGTC -3’ (reverse)

GAPDH: see above

### Analysis of N4BP1 expression in clinical liver samples

To assess the N4BP1 expression of HBV-HCC resection samples, we confirmed that all specimens were HBs-Ag (+), HBe-Ag (-), and HBe-Ab (+) as documented in the patient medical records. Based on HBV DNA data in the serum, the samples were divided into an HBV DNA positive group (n=9) and an HBV DNA negative group (n=8). In all cases, RNA was extracted from the non-cancerous areas of resected tissues using the RNeasy Mini Kit and *N4BP1* and *ISG15* mRNA expression were measured by qPCR. This study was approved by the Institutional Review Board of the Graduate School of Medicine, Hokkaido University (No. 14-015, 022-0195). All patients participating in this study provided written informed consent for the use of their samples and clinical data and they were informed of the opportunity to opt out.

### Northern blotting

Huh7 cells were co-transfected with a plasmid containing the 1.3-fold-overlength genome of HBV genotype C and N4BP1 using TransIT LT-1 transfection reagent, and cells were collected at 3 d.p.t.. Total RNA was extracted using Isogen (Nippon Gene, Tokyo, Japan) and subjected to electrophoresis in 1% agarose-formaldehyde gel and morpholinepropanesulfonic acid buffer (Nacalai Tesque). Ribosomal RNA was visualized by ethidium bromide staining, and electrophoresed RNA was transferred onto a positively charged nylon membrane (Roche, Basel-Stadt, Switzerland). An HBV RNA probe and GAPDH RNA probe were linearized and synthesized by digoxigenin RNA labeling kit (Roche). Hybridization and detection were performed using the DIG Northern Starter kit (Roche) following the manufacturer’s protocol. The signals were detected by LuminoGraph I (Atto).

### RNA Immunoprecipitation (RIP) assay

Huh7 cells were co-transfected with a plasmid containing the 1.3-fold-overlength genome of HBV genotype C and HA-tagged N4BP1 wildtype or domain deficient mutants or HA control vector using TransIT LT-1 transfection reagent, and cells were collected at 3 d.p.t. Cells were collected and immunoprecipitated with anti-HA-tag monoclonal antibody magnetic beads (Medical & Biological Laboratories Co., Ltd., Tokyo, Japan). Immunoprecipitation and RNA extraction were performed using RiboCluster Profiler RIP-Assay Kit (Medical & Biological Laboratories Co., Ltd., Tokyo, Japan) following the manufacture’s protocol. Western blotting and pgRNA qPCR were measured as describe above.

### HBeAg CLIA, HBs Ag ELISA and HBcrAg ELISA

N4BP1-overexpressing PHHs were infected with HBV genotype C (5 Geq/cell). At 14 d.p.i., RNA was extracted from the cells and pgRNA was measured as described above. HBV proteins in the supernatant were assessed as follows: HBeAg CLIA (Ig Biotechnology, Burlingame, USA); HBs S Antigen Quantitative ELISA Kit, Rapid-II (Takara Bio) following manufacturers’ protocol; and HBcr Ag assay (42) by SRL, Inc. (Tokyo, Japan).

### RNA sequencing

PHHs were analyzed for RNA sequencing. N4BP1-overexpressing PHHs were infected with HBV genotype C (5 Geq/cell) as described above. At 3 d.p.i., total RNA in PHHs was extracted using RNeasy Mini Kit following the manufacturer’s protocol. RNA sequencing was performed by Osaka University (Genome Information Research Center, Research Institute for Microbial Diseases, Osaka University, Osaka, Japan). We ran 2 well (N4BP1-overexpressing) vs 2 well (control) samples. Whole transcriptome sequencing was applied to RNA samples using the Illumina NovaSeq 6000 platform in a 101 bp single-end mode. Sequenced reads were mapped to the human reference genome sequence (hg19) using TopHat version 2.2.1 in combination with Bowtie 2 version 2.2.3 and SAMtools version 1.0. The number of fragments per kilobase of exon per million mapped fragments was calculated using Cufflinks version 2.2.1. Following the exclusion of genes with expression counts below 50, differential expression analysis was conducted utilizing the DESeq2 package (version 1.42.0) (43). Genes were considered significantly differentially expressed if they satisfied the criteria of an adjusted P-value less than 0.1 and an absolute fold change greater than 2. Subsequently, pathway enrichment analysis of the differentially expressed genes was performed using the Kyoto Encyclopedia of Genes and Genomes database (44) with the clusterProfiler package (version 4.10.0) (45). Plots were generated using the ggplot2 package (version 3.5.1) (46). To review GEO data, access https://www.ncbi.nlm.nih.gov/geo/query/acc.cgi?acc=GSE225322. The accession No. is GSE225322.

### Statistical analysis

Data are averages of three or more independent experiments. All data in figures are presented as the mean ± standard deviation (SD). Evaluation of data was performed by Mann-Whitney *U* test. *P*-value of < 0.05 was considered statistically significant. Statistical analysis was performed using JMP Pro version 16.

## Abbreviations

cccDNA: covalently closed circular DNA
d.p.i.: days post-infection
d.p.t.: days post-transfection
ETV: entecavir
HBV: hepatitis B virus
HBcrAg: hepatitis B core-related antigens
HCC: hepatocellular carcinoma
IFN: interferon
RIP assay: RNA Immunoprecipitation assay
ISG: interferon-stimulated gene
KO: knockout
MALT1: mucosa-associated lymphoid tissue lymphoma translocation 1
NA: nucleos(t)ide analog
NTCP: sodium-taurocholate cotransporting polypeptide
N4BP1: NEDD4-binding protein 1
pgRNA: pregenomic RNA
PHH: primary human hepatocyte
qPCR: quantitative PCR
RBP: RNA-binding protein
rcDNA: relaxed circular DNA
WB: western blotting
WT: wild type

## Data Availability

The GEO data that support the findings of this study are openly available in https://www.ncbi.nlm.nih.gov/geo/query/acc.cgi?acc=GSE225322. The accession No. is GSE225322.

## Acknowledgements

We thank Haruko Kubo, Mizuho Tetsuka, and Saya Shimamura for their secretarial work and Mikiko Ishibashi, Minako Hommura and Hiroko Azuma for their technical assistance. We are grateful to Takaji Wakita (NIID) for providing HepG2-hNTCP C4 cells and HepAD38.7 cells. We also thank Jenna Gaska for English proofreading.

This work was supported in part by the Japan Agency for Medical Research and Development (AMED) under Grant Numbers 22fk0310501h0001, 22fk0310506h0001, 22fk0310513h0001, 22fk0310524h0001, and 22fk0310523h0001; AMED CREST under Grant Number JP22gm1610008; JSPS KAKENHI Fund for the Promotion of Joint International Research (International Leading Research) under grant number JP23K20041.

Conceptualization, Y. M., O. T., A. T., and T. F.; Data curation, N. K., S. S., R. S., T. T., D. O., and T. F.; Formal analysis, N. K., S. S., R. S., T. T., D. O., K. M., and T. F.; Funding acquisition, T. F.; Investigation, N. K., S. S., Y. S., T. I., K. N., D. O., K. M., L. L., Y. K., and Y. K.; Methodology, N. K., S. S., R. S., T. T., S. H., Y. T., A. T., and T. F.; Project administration, A. T. and T. F.; Resources, D. O., Y. K., Y. M., O. T., A. T., and T. F.; Software, D. O.; Supervision, A. T., T. F.; Validation, N. K., S. S., Y. S., T. S., T. I., and K. N.; Visualization, N. K., S. S., and T. F.; Writing - original draft, N. K., S. S., D. O., Y. K., K. M., and T. F.; Writing - review & editing, N. K., S. S., R. S., T. T., Y. S., T. S., T. I., K. N., D. O., Y. K., Y. M., O. T., A. T., and T. F. All authors have approved the submitted version of the manuscript and agreed to be accountable for any part of the work.

