## Supplementary material for "NEDD4-binding protein 1 suppresses HBV replication by degrading HBV RNAs": Table S1 to S2

**S1 Table　Differential gene expression profile by RNA-seq of HBV-infected PHHs overexpressing N4BP1 compared to HBV-infected control PHHs.**

| Factor name | baseMean | log2FoldChange | lfcSE | stat | pvalue | padj | distance | significance |
| --- | --- | --- | --- | --- | --- | --- | --- | --- |
| N4BP1 | 11471.14 | 4.223 | 0.081 | 51.88 | 0.00E+00 | 0.00E+00 | Inf | Upregulated |
| NNMT | 1189.55 | 3.009 | 0.137 | 21.94 | 1.08E-106 | 6.59E-103 | 102.226 | Upregulated |
| FGB | 70363.60 | 1.589 | 0.073 | 21.79 | 3.14E-105 | 1.28E-101 | 100.905 | Upregulated |
| KRT8 | 27810.42 | 1.444 | 0.071 | 20.27 | 2.14E-91 | 6.55E-88 | 87.196 | Upregulated |
| FSTL1 | 2342.94 | 1.859 | 0.097 | 19.13 | 1.34E-81 | 3.27E-78 | 77.507 | Upregulated |
| KRT18 | 11689.26 | 1.372 | 0.078 | 17.48 | 2.03E-68 | 4.14E-65 | 64.398 | Upregulated |
| FGG | 74088.49 | 1.262 | 0.075 | 16.84 | 1.15E-63 | 2.01E-60 | 59.710 | Upregulated |
| SAA2 | 811.77 | 2.184 | 0.132 | 16.55 | 1.49E-61 | 2.02E-58 | 57.736 | Upregulated |
| SAA1 | 4684.75 | 1.545 | 0.100 | 15.53 | 2.19E-54 | 2.43E-51 | 50.638 | Upregulated |
| ANGPTL4 | 6260.94 | 1.639 | 0.106 | 15.48 | 4.95E-54 | 5.04E-51 | 50.324 | Upregulated |
| MT1E | 441.04 | 2.482 | 0.163 | 15.27 | 1.22E-52 | 1.15E-49 | 49.002 | Upregulated |
| CCND1 | 1089.74 | 1.818 | 0.123 | 14.76 | 2.57E-49 | 2.24E-46 | 45.686 | Upregulated |
| FGA | 120757.14 | 1.129 | 0.079 | 14.34 | 1.27E-46 | 9.15E-44 | 43.053 | Upregulated |
| BCL2L1 | 2946.48 | 1.145 | 0.085 | 13.49 | 1.70E-41 | 1.16E-38 | 37.954 | Upregulated |
| KRT7 | 871.67 | 3.792 | 0.286 | 13.25 | 4.38E-40 | 2.81E-37 | 36.747 | Upregulated |
| ID1 | 2106.95 | 1.333 | 0.101 | 13.21 | 7.19E-40 | 4.39E-37 | 36.382 | Upregulated |
| EPAS1 | 3080.23 | 1.151 | 0.088 | 13.10 | 3.22E-39 | 1.87E-36 | 35.746 | Upregulated |
| CCND2 | 520.95 | 1.809 | 0.145 | 12.48 | 9.81E-36 | 5.00E-33 | 32.352 | Upregulated |
| TRNP1 | 1915.79 | 1.271 | 0.103 | 12.36 | 4.06E-35 | 1.98E-32 | 31.728 | Upregulated |
| C10orf54 | 611.04 | 1.596 | 0.133 | 12.02 | 2.74E-33 | 1.29E-30 | 29.933 | Upregulated |
| SYT12 | 324.54 | 2.271 | 0.190 | 11.92 | 9.36E-33 | 4.09E-30 | 29.476 | Upregulated |
| AHNAK2 | 226.50 | 2.465 | 0.215 | 11.49 | 1.43E-30 | 5.15E-28 | 27.399 | Upregulated |
| SLC2A2 | 2432.96 | 1.233 | 0.108 | 11.45 | 2.45E-30 | 8.56E-28 | 27.095 | Upregulated |
| SLC35C1 | 2293.29 | 1.037 | 0.093 | 11.18 | 5.01E-29 | 1.70E-26 | 25.790 | Upregulated |
| ANXA3 | 372.44 | 2.157 | 0.194 | 11.12 | 9.99E-29 | 3.21E-26 | 25.584 | Upregulated |
| NBEAL2 | 597.84 | 1.550 | 0.140 | 11.09 | 1.35E-28 | 4.23E-26 | 25.421 | Upregulated |
| S100A11 | 1740.92 | 1.150 | 0.108 | 10.69 | 1.08E-26 | 2.88E-24 | 23.569 | Upregulated |
| MT2A | 1497.14 | 2.668 | 0.250 | 10.66 | 1.60E-26 | 4.17E-24 | 23.532 | Upregulated |
| NR0B2 | 164.15 | 2.731 | 0.258 | 10.58 | 3.70E-26 | 9.18E-24 | 23.199 | Upregulated |
| SLC7A7 | 413.42 | 1.635 | 0.155 | 10.58 | 3.75E-26 | 9.18E-24 | 23.095 | Upregulated |
| CPA4 | 137.93 | 2.983 | 0.290 | 10.27 | 9.64E-25 | 1.90E-22 | 21.925 | Upregulated |
| GALK1 | 964.70 | 1.216 | 0.118 | 10.29 | 8.10E-25 | 1.65E-22 | 21.816 | Upregulated |
| MYL9 | 303.84 | 2.131 | 0.208 | 10.24 | 1.33E-24 | 2.49E-22 | 21.708 | Upregulated |
| DUSP6 | 275.19 | 1.954 | 0.193 | 10.11 | 4.84E-24 | 8.84E-22 | 21.144 | Upregulated |
| MCAM | 494.51 | 1.515 | 0.154 | 9.83 | 8.21E-23 | 1.30E-20 | 19.942 | Upregulated |
| TAT | 3852.07 | 1.038 | 0.106 | 9.81 | 1.00E-22 | 1.57E-20 | 19.832 | Upregulated |
| ABALON | 1090.24 | 1.040 | 0.107 | 9.68 | 3.58E-22 | 5.40E-20 | 19.295 | Upregulated |
| CUX2 | 559.43 | 1.362 | 0.142 | 9.62 | 6.81E-22 | 9.79E-20 | 19.058 | Upregulated |
| EPHA2 | 922.79 | 1.075 | 0.113 | 9.51 | 1.89E-21 | 2.65E-19 | 18.608 | Upregulated |
| MFSD2A | 185.09 | 2.400 | 0.254 | 9.44 | 3.57E-21 | 4.96E-19 | 18.462 | Upregulated |
| MT1X | 162.38 | 2.511 | 0.269 | 9.33 | 1.06E-20 | 1.43E-18 | 18.022 | Upregulated |
| LOXL4 | 660.17 | 1.445 | 0.155 | 9.32 | 1.19E-20 | 1.58E-18 | 17.861 | Upregulated |
| TAGLN | 208.81 | 2.002 | 0.221 | 9.06 | 1.30E-19 | 1.63E-17 | 16.906 | Upregulated |
| TGM2 | 3127.81 | 2.409 | 0.268 | 8.99 | 2.37E-19 | 2.92E-17 | 16.709 | Upregulated |
| SDC4 | 2311.64 | 1.028 | 0.115 | 8.95 | 3.54E-19 | 4.28E-17 | 16.401 | Upregulated |
| NAT8 | 402.48 | 1.372 | 0.155 | 8.85 | 8.88E-19 | 1.05E-16 | 16.036 | Upregulated |
| SLC25A25 | 396.24 | 1.399 | 0.158 | 8.84 | 9.94E-19 | 1.17E-16 | 15.994 | Upregulated |
| UNC5CL | 429.19 | 1.305 | 0.155 | 8.44 | 3.29E-17 | 3.47E-15 | 14.519 | Upregulated |
| ETV1 | 291.18 | 1.503 | 0.178 | 8.42 | 3.79E-17 | 3.93E-15 | 14.484 | Upregulated |
| CNN2 | 908.97 | 1.197 | 0.144 | 8.29 | 1.09E-16 | 1.08E-14 | 14.018 | Upregulated |
| IL18 | 136.71 | 2.384 | 0.289 | 8.24 | 1.68E-16 | 1.61E-14 | 13.997 | Upregulated |
| ETV5 | 105.02 | 2.531 | 0.314 | 8.05 | 8.01E-16 | 7.14E-14 | 13.388 | Upregulated |
| TGFB2 | 165.15 | 2.003 | 0.255 | 7.85 | 4.30E-15 | 3.58E-13 | 12.606 | Upregulated |
| MT1G | 134.20 | 2.275 | 0.300 | 7.59 | 3.28E-14 | 2.54E-12 | 11.817 | Upregulated |
| MYOF | 456.37 | 1.170 | 0.155 | 7.56 | 3.88E-14 | 2.93E-12 | 11.593 | Upregulated |
| SDS | 47.14 | 4.159 | 0.581 | 7.16 | 7.86E-13 | 4.98E-11 | 11.111 | Upregulated |
| SPRY4 | 75.58 | 2.673 | 0.367 | 7.29 | 3.09E-13 | 2.07E-11 | 11.014 | Upregulated |
| FBLIM1 | 283.70 | 1.294 | 0.179 | 7.21 | 5.52E-13 | 3.57E-11 | 10.527 | Upregulated |
| SLC16A12 | 214.82 | 1.540 | 0.221 | 6.98 | 2.95E-12 | 1.68E-10 | 9.895 | Upregulated |
| CRP | 80.82 | 2.486 | 0.363 | 6.85 | 7.28E-12 | 3.94E-10 | 9.728 | Upregulated |
| FLNC | 174.91 | 1.541 | 0.223 | 6.92 | 4.47E-12 | 2.49E-10 | 9.726 | Upregulated |
| RBMS2 | 347.10 | 1.124 | 0.163 | 6.91 | 4.89E-12 | 2.69E-10 | 9.636 | Upregulated |
| DEFB1 | 431.44 | 1.021 | 0.152 | 6.71 | 1.98E-11 | 1.01E-09 | 9.055 | Upregulated |
| THBS1 | 348.31 | 1.283 | 0.193 | 6.65 | 3.01E-11 | 1.46E-09 | 8.930 | Upregulated |
| ATP2B4 | 550.46 | 1.016 | 0.155 | 6.54 | 6.00E-11 | 2.74E-09 | 8.623 | Upregulated |
| SH3YL1 | 385.54 | 1.027 | 0.158 | 6.49 | 8.72E-11 | 3.87E-09 | 8.474 | Upregulated |
| RNF157 | 97.55 | 1.911 | 0.298 | 6.40 | 1.51E-10 | 6.43E-09 | 8.412 | Upregulated |
| CCAT1 | 62.61 | 2.672 | 0.427 | 6.26 | 3.77E-10 | 1.53E-08 | 8.259 | Upregulated |
| PRICKLE2 | 216.34 | 1.221 | 0.200 | 6.10 | 1.08E-09 | 4.13E-08 | 7.484 | Upregulated |
| KRT80 | 29.18 | 4.207 | 0.753 | 5.59 | 2.31E-08 | 6.78E-07 | 7.467 | Upregulated |
| LBH | 123.98 | 1.881 | 0.315 | 5.97 | 2.32E-09 | 8.28E-08 | 7.328 | Upregulated |
| ATOH8 | 256.52 | 1.110 | 0.185 | 6.02 | 1.77E-09 | 6.45E-08 | 7.276 | Upregulated |
| FAM105A | 94.06 | 1.734 | 0.301 | 5.77 | 8.05E-09 | 2.60E-07 | 6.809 | Upregulated |
| C9 | 102.29 | 1.668 | 0.291 | 5.73 | 1.03E-08 | 3.25E-07 | 6.699 | Upregulated |
| SVEP1 | 67.11 | 2.093 | 0.370 | 5.65 | 1.58E-08 | 4.80E-07 | 6.656 | Upregulated |
| PMEPA1 | 92.30 | 1.973 | 0.353 | 5.60 | 2.20E-08 | 6.48E-07 | 6.496 | Upregulated |
| PDE11A | 122.01 | 1.478 | 0.265 | 5.57 | 2.51E-08 | 7.33E-07 | 6.311 | Upregulated |
| LINC01488 | 91.81 | 1.755 | 0.320 | 5.48 | 4.21E-08 | 1.18E-06 | 6.183 | Upregulated |
| UBASH3B | 64.86 | 2.178 | 0.404 | 5.39 | 7.02E-08 | 1.87E-06 | 6.129 | Upregulated |
| BTBD6 | 273.76 | 1.031 | 0.189 | 5.46 | 4.74E-08 | 1.31E-06 | 5.974 | Upregulated |
| KNDC1 | 27.44 | 3.319 | 0.660 | 5.03 | 4.97E-07 | 1.10E-05 | 5.969 | Upregulated |
| BCO2 | 186.65 | 1.142 | 0.212 | 5.38 | 7.35E-08 | 1.94E-06 | 5.824 | Upregulated |
| TMEM147 | 217.10 | 1.065 | 0.198 | 5.38 | 7.27E-08 | 1.93E-06 | 5.814 | Upregulated |
| MMD | 154.96 | 1.245 | 0.234 | 5.33 | 1.00E-07 | 2.55E-06 | 5.731 | Upregulated |
| EFHD1 | 171.44 | 1.161 | 0.229 | 5.08 | 3.77E-07 | 8.61E-06 | 5.197 | Upregulated |
| MMP24 | 107.54 | 1.405 | 0.282 | 4.99 | 6.14E-07 | 1.32E-05 | 5.077 | Upregulated |
| HAMP | 3098.72 | 1.062 | 0.213 | 4.99 | 5.95E-07 | 1.29E-05 | 5.004 | Upregulated |
| MT1L | 68.11 | 1.734 | 0.361 | 4.81 | 1.55E-06 | 3.01E-05 | 4.843 | Upregulated |
| COTL1 | 89.32 | 1.466 | 0.302 | 4.85 | 1.23E-06 | 2.45E-05 | 4.838 | Upregulated |
| BTBD11 | 205.61 | 1.066 | 0.217 | 4.91 | 9.34E-07 | 1.94E-05 | 4.832 | Upregulated |
| UBALD2 | 139.49 | 1.241 | 0.254 | 4.88 | 1.06E-06 | 2.16E-05 | 4.828 | Upregulated |
| SLC25A18 | 97.07 | 1.433 | 0.297 | 4.83 | 1.38E-06 | 2.72E-05 | 4.786 | Upregulated |
| IP6K3 | 102.80 | 1.332 | 0.286 | 4.66 | 3.23E-06 | 5.80E-05 | 4.441 | Upregulated |
| ITGB6 | 117.77 | 1.287 | 0.279 | 4.62 | 3.90E-06 | 6.78E-05 | 4.363 | Upregulated |
| LINC00152 | 96.06 | 1.393 | 0.307 | 4.54 | 5.68E-06 | 9.58E-05 | 4.253 | Upregulated |
| TSPAN15 | 113.04 | 1.218 | 0.269 | 4.52 | 6.05E-06 | 1.02E-04 | 4.175 | Upregulated |
| FGFR1 | 163.44 | 1.081 | 0.238 | 4.54 | 5.51E-06 | 9.37E-05 | 4.171 | Upregulated |
| KDELR3 | 47.65 | 1.869 | 0.428 | 4.37 | 1.25E-05 | 1.96E-04 | 4.152 | Upregulated |
| MT1F | 27.19 | 2.434 | 0.622 | 3.92 | 9.02E-05 | 1.08E-03 | 3.836 | Upregulated |
| UNC5B | 91.33 | 1.369 | 0.321 | 4.27 | 1.99E-05 | 2.93E-04 | 3.790 | Upregulated |
| VCAN | 78.84 | 1.463 | 0.348 | 4.20 | 2.67E-05 | 3.78E-04 | 3.722 | Upregulated |
| ICOSLG | 118.78 | 1.117 | 0.264 | 4.24 | 2.25E-05 | 3.24E-04 | 3.664 | Upregulated |
| FNDC5 | 63.38 | 1.537 | 0.375 | 4.10 | 4.20E-05 | 5.62E-04 | 3.596 | Upregulated |
| DNAH5 | 79.09 | 1.400 | 0.340 | 4.12 | 3.86E-05 | 5.20E-04 | 3.570 | Upregulated |
| PLSCR3 | 82.33 | 1.480 | 0.362 | 4.09 | 4.30E-05 | 5.74E-04 | 3.563 | Upregulated |
| NEXN | 103.30 | 1.190 | 0.287 | 4.15 | 3.38E-05 | 4.64E-04 | 3.539 | Upregulated |
| AKAP12 | 1086.42 | 1.076 | 0.261 | 4.12 | 3.73E-05 | 5.06E-04 | 3.467 | Upregulated |
| TSPAN18 | 50.89 | 1.620 | 0.407 | 3.98 | 6.95E-05 | 8.66E-04 | 3.465 | Upregulated |
| CMTM3 | 94.32 | 1.173 | 0.291 | 4.03 | 5.54E-05 | 7.09E-04 | 3.361 | Upregulated |
| ACTA1 | 31.44 | 2.012 | 0.539 | 3.73 | 1.91E-04 | 2.08E-03 | 3.353 | Upregulated |
| ABLIM2 | 33.93 | 1.881 | 0.500 | 3.76 | 1.70E-04 | 1.88E-03 | 3.311 | Upregulated |
| MYADM | 132.00 | 1.026 | 0.255 | 4.03 | 5.68E-05 | 7.26E-04 | 3.303 | Upregulated |
| TPM4 | 2732.49 | 1.054 | 0.264 | 4.00 | 6.41E-05 | 8.10E-04 | 3.266 | Upregulated |
| S100A9 | 75.85 | 1.293 | 0.331 | 3.90 | 9.45E-05 | 1.13E-03 | 3.218 | Upregulated |
| C2orf88 | 114.79 | 1.045 | 0.264 | 3.96 | 7.50E-05 | 9.24E-04 | 3.209 | Upregulated |
| TEAD4 | 63.63 | 1.371 | 0.356 | 3.85 | 1.16E-04 | 1.35E-03 | 3.180 | Upregulated |
| TMEM54 | 107.55 | 1.053 | 0.273 | 3.86 | 1.15E-04 | 1.34E-03 | 3.059 | Upregulated |
| FSCN1 | 98.81 | 1.148 | 0.303 | 3.79 | 1.52E-04 | 1.71E-03 | 2.997 | Upregulated |
| CCDC113 | 32.43 | 1.798 | 0.518 | 3.47 | 5.15E-04 | 4.81E-03 | 2.933 | Upregulated |
| CDR2L | 66.87 | 1.308 | 0.355 | 3.68 | 2.33E-04 | 2.49E-03 | 2.914 | Upregulated |
| MAGI2-AS3 | 71.61 | 1.222 | 0.333 | 3.67 | 2.39E-04 | 2.54E-03 | 2.868 | Upregulated |
| SOBP | 97.56 | 1.087 | 0.294 | 3.70 | 2.16E-04 | 2.32E-03 | 2.850 | Upregulated |
| TMOD2 | 79.35 | 1.186 | 0.326 | 3.63 | 2.81E-04 | 2.90E-03 | 2.801 | Upregulated |
| GRAMD1B | 71.87 | 1.205 | 0.334 | 3.61 | 3.11E-04 | 3.15E-03 | 2.777 | Upregulated |
| SDC3 | 100.81 | 1.041 | 0.286 | 3.63 | 2.81E-04 | 2.90E-03 | 2.742 | Upregulated |
| TESC | 62.88 | 1.263 | 0.364 | 3.48 | 5.10E-04 | 4.77E-03 | 2.643 | Upregulated |
| TMEM154 | 49.41 | 1.376 | 0.403 | 3.41 | 6.46E-04 | 5.78E-03 | 2.627 | Upregulated |
| WSCD1 | 75.10 | 1.159 | 0.335 | 3.46 | 5.49E-04 | 5.07E-03 | 2.571 | Upregulated |
| GDPD5 | 36.42 | 1.556 | 0.488 | 3.19 | 1.44E-03 | 1.11E-02 | 2.497 | Upregulated |
| PDGFB | 29.69 | 1.631 | 0.525 | 3.11 | 1.88E-03 | 1.38E-02 | 2.474 | Upregulated |
| TBX15 | 66.37 | 1.189 | 0.356 | 3.34 | 8.42E-04 | 7.16E-03 | 2.453 | Upregulated |
| TMEM231 | 27.19 | 1.672 | 0.549 | 3.05 | 2.32E-03 | 1.64E-02 | 2.447 | Upregulated |
| GMPR | 88.08 | 1.028 | 0.303 | 3.39 | 7.00E-04 | 6.17E-03 | 2.437 | Upregulated |
| ASPG | 28.44 | 1.616 | 0.531 | 3.04 | 2.33E-03 | 1.64E-02 | 2.409 | Upregulated |
| SYTL2 | 37.93 | 1.434 | 0.458 | 3.13 | 1.75E-03 | 1.30E-02 | 2.368 | Upregulated |
| HID1 | 77.61 | 1.053 | 0.318 | 3.31 | 9.40E-04 | 7.83E-03 | 2.355 | Upregulated |
| NT5DC2 | 40.17 | 1.408 | 0.454 | 3.10 | 1.93E-03 | 1.41E-02 | 2.325 | Upregulated |
| NPM3 | 59.14 | 1.194 | 0.371 | 3.22 | 1.30E-03 | 1.02E-02 | 2.320 | Upregulated |
| SFN | 27.94 | 1.582 | 0.538 | 2.94 | 3.26E-03 | 2.14E-02 | 2.300 | Upregulated |
| GLIPR2 | 52.65 | 1.229 | 0.395 | 3.12 | 1.83E-03 | 1.35E-02 | 2.237 | Upregulated |
| CHST3 | 53.65 | 1.204 | 0.385 | 3.13 | 1.76E-03 | 1.31E-02 | 2.235 | Upregulated |
| MAP3K6 | 72.61 | 1.042 | 0.328 | 3.18 | 1.47E-03 | 1.13E-02 | 2.207 | Upregulated |
| TIAM2 | 51.90 | 1.200 | 0.387 | 3.10 | 1.94E-03 | 1.42E-02 | 2.203 | Upregulated |
| NPM2 | 53.65 | 1.172 | 0.393 | 2.98 | 2.84E-03 | 1.91E-02 | 2.080 | Upregulated |
| GLIPR1 | 40.92 | 1.270 | 0.435 | 2.92 | 3.52E-03 | 2.29E-02 | 2.075 | Upregulated |
| NCF2 | 27.44 | 1.479 | 0.540 | 2.74 | 6.14E-03 | 3.53E-02 | 2.073 | Upregulated |
| FGF19 | 47.41 | 1.220 | 0.415 | 2.94 | 3.29E-03 | 2.16E-02 | 2.065 | Upregulated |
| LOC100126784 | 44.16 | 1.263 | 0.441 | 2.86 | 4.19E-03 | 2.62E-02 | 2.024 | Upregulated |
| GLP2R | 62.88 | 1.049 | 0.355 | 2.95 | 3.13E-03 | 2.07E-02 | 1.984 | Upregulated |
| HES2 | 59.89 | 1.052 | 0.360 | 2.92 | 3.47E-03 | 2.26E-02 | 1.954 | Upregulated |
| EPHB6 | 62.39 | 1.032 | 0.353 | 2.93 | 3.44E-03 | 2.24E-02 | 1.946 | Upregulated |
| ULBP2 | 39.43 | 1.237 | 0.445 | 2.78 | 5.39E-03 | 3.18E-02 | 1.943 | Upregulated |
| SAMD11 | 34.94 | 1.319 | 0.488 | 2.71 | 6.82E-03 | 3.82E-02 | 1.937 | Upregulated |
| SLC19A1 | 42.67 | 1.192 | 0.425 | 2.80 | 5.04E-03 | 3.02E-02 | 1.931 | Upregulated |
| CXCL8 | 128.73 | 1.466 | 0.585 | 2.50 | 1.23E-02 | 6.01E-02 | 1.907 | Upregulated |
| PCNXL2 | 40.43 | 1.203 | 0.440 | 2.73 | 6.29E-03 | 3.59E-02 | 1.880 | Upregulated |
| SLC4A11 | 35.19 | 1.283 | 0.483 | 2.66 | 7.82E-03 | 4.26E-02 | 1.878 | Upregulated |
| ARID3A | 40.67 | 1.216 | 0.447 | 2.72 | 6.59E-03 | 3.71E-02 | 1.877 | Upregulated |
| PTRH1 | 34.18 | 1.275 | 0.479 | 2.66 | 7.81E-03 | 4.26E-02 | 1.872 | Upregulated |
| FAM86EP | 36.68 | 1.224 | 0.458 | 2.67 | 7.47E-03 | 4.11E-02 | 1.849 | Upregulated |
| NRP2 | 25.95 | 1.368 | 0.547 | 2.50 | 1.23E-02 | 6.02E-02 | 1.833 | Upregulated |
| ZWILCH | 53.65 | 1.079 | 0.391 | 2.76 | 5.81E-03 | 3.38E-02 | 1.824 | Upregulated |
| RTN4RL1 | 28.19 | 1.337 | 0.530 | 2.52 | 1.16E-02 | 5.75E-02 | 1.824 | Upregulated |
| DUSP18 | 46.66 | 1.115 | 0.410 | 2.72 | 6.57E-03 | 3.70E-02 | 1.814 | Upregulated |
| FGF21 | 27.44 | 1.347 | 0.552 | 2.44 | 1.47E-02 | 6.86E-02 | 1.780 | Upregulated |
| FAM129A | 39.18 | 1.180 | 0.452 | 2.61 | 8.96E-03 | 4.72E-02 | 1.775 | Upregulated |
| C1orf111 | 38.18 | 1.171 | 0.458 | 2.55 | 1.06E-02 | 5.38E-02 | 1.727 | Upregulated |
| IQCA1 | 36.68 | 1.178 | 0.465 | 2.53 | 1.14E-02 | 5.67E-02 | 1.715 | Upregulated |
| NAV1 | 43.42 | 1.111 | 0.429 | 2.59 | 9.55E-03 | 4.95E-02 | 1.714 | Upregulated |
| TIGD3 | 26.20 | 1.253 | 0.540 | 2.32 | 2.04E-02 | 8.74E-02 | 1.640 | Upregulated |
| CHST15 | 40.93 | 1.104 | 0.445 | 2.48 | 1.31E-02 | 6.28E-02 | 1.633 | Upregulated |
| MAP4K2 | 39.93 | 1.093 | 0.439 | 2.49 | 1.29E-02 | 6.20E-02 | 1.629 | Upregulated |
| TCEAL3 | 33.19 | 1.163 | 0.483 | 2.41 | 1.60E-02 | 7.29E-02 | 1.627 | Upregulated |
| NPL | 29.19 | 1.167 | 0.514 | 2.27 | 2.31E-02 | 9.56E-02 | 1.550 | Upregulated |
| CERCAM | 42.67 | 1.035 | 0.431 | 2.41 | 1.62E-02 | 7.34E-02 | 1.536 | Upregulated |
| FICD | 42.93 | 1.048 | 0.438 | 2.39 | 1.68E-02 | 7.55E-02 | 1.535 | Upregulated |
| IGFLR1 | 33.69 | 1.095 | 0.477 | 2.29 | 2.18E-02 | 9.17E-02 | 1.508 | Upregulated |
| ANKRD1 | 2465.17 | 1.030 | 0.446 | 2.31 | 2.10E-02 | 8.91E-02 | 1.471 | Upregulated |

| Factor name | baseMean | log2FoldChange | lfcSE | stat | pvalue | padj | distance | significance |
| --- | --- | --- | --- | --- | --- | --- | --- | --- |
| COL6A2 | 312.77 | -3.593 | 0.215 | -16.70 | 1.28E-62 | 1.95E-59 | 58.819 | Downregulated |
| ITIH3 | 4272.13 | -1.445 | 0.093 | -15.61 | 6.23E-55 | 7.62E-52 | 51.139 | Downregulated |
| FADS1 | 12316.26 | -1.278 | 0.087 | -14.75 | 3.00E-49 | 2.44E-46 | 45.630 | Downregulated |
| TTR | 45920.62 | -1.090 | 0.074 | -14.73 | 3.86E-49 | 2.95E-46 | 45.543 | Downregulated |
| CYP4F2 | 1809.39 | -1.202 | 0.095 | -12.71 | 4.97E-37 | 2.76E-34 | 33.580 | Downregulated |
| FER1L4 | 880.33 | -1.764 | 0.148 | -11.91 | 1.03E-32 | 4.34E-30 | 29.415 | Downregulated |
| UCHL1 | 683.26 | -1.499 | 0.128 | -11.75 | 6.85E-32 | 2.70E-29 | 28.608 | Downregulated |
| MAP3K14 | 890.46 | -1.295 | 0.118 | -10.97 | 5.48E-28 | 1.67E-25 | 24.810 | Downregulated |
| BIRC3 | 2340.63 | -1.313 | 0.122 | -10.76 | 5.10E-27 | 1.45E-24 | 23.875 | Downregulated |
| MTRNR2L2 | 4266.49 | -1.015 | 0.095 | -10.72 | 8.39E-27 | 2.28E-24 | 23.664 | Downregulated |
| NQO1 | 2789.02 | -1.091 | 0.103 | -10.60 | 3.13E-26 | 7.98E-24 | 23.124 | Downregulated |
| PNN | 1244.09 | -1.190 | 0.113 | -10.55 | 4.99E-26 | 1.17E-23 | 22.962 | Downregulated |
| MTRNR2L8 | 2408.68 | -1.052 | 0.101 | -10.46 | 1.36E-25 | 3.08E-23 | 22.536 | Downregulated |
| MTRNR2L9 | 2479.24 | -1.161 | 0.111 | -10.44 | 1.60E-25 | 3.55E-23 | 22.480 | Downregulated |
| SLCO1B1 | 488.95 | -1.523 | 0.147 | -10.38 | 2.94E-25 | 6.31E-23 | 22.252 | Downregulated |
| XPNPEP2 | 453.75 | -1.534 | 0.149 | -10.31 | 6.04E-25 | 1.27E-22 | 21.949 | Downregulated |
| COL6A1 | 741.22 | -1.461 | 0.142 | -10.30 | 7.09E-25 | 1.47E-22 | 21.882 | Downregulated |
| ALDH1L1 | 3073.16 | -1.029 | 0.100 | -10.25 | 1.19E-24 | 2.29E-22 | 21.664 | Downregulated |
| CYP2D6 | 462.52 | -1.588 | 0.160 | -9.94 | 2.74E-23 | 4.65E-21 | 20.395 | Downregulated |
| CYP1A2 | 412.79 | -1.488 | 0.153 | -9.70 | 3.13E-22 | 4.78E-20 | 19.378 | Downregulated |
| COL7A1 | 308.46 | -1.766 | 0.201 | -8.81 | 1.29E-18 | 1.50E-16 | 15.921 | Downregulated |
| CYP7A1 | 68.45 | -4.725 | 0.548 | -8.63 | 6.33E-18 | 7.23E-16 | 15.861 | Downregulated |
| VCAM1 | 78.94 | -3.365 | 0.391 | -8.60 | 8.16E-18 | 8.98E-16 | 15.418 | Downregulated |
| PDE4C | 235.76 | -1.889 | 0.221 | -8.53 | 1.49E-17 | 1.62E-15 | 14.910 | Downregulated |
| NUDT4P2 | 136.42 | -10.538 | 1.459 | -7.22 | 5.19E-13 | 3.39E-11 | 14.855 | Downregulated |
| ADH1A | 474.00 | -1.378 | 0.162 | -8.52 | 1.55E-17 | 1.67E-15 | 14.841 | Downregulated |
| VAMP1 | 457.53 | -1.464 | 0.174 | -8.43 | 3.42E-17 | 3.57E-15 | 14.521 | Downregulated |
| MIR210HG | 289.92 | -1.684 | 0.200 | -8.41 | 4.21E-17 | 4.32E-15 | 14.462 | Downregulated |
| ZBED6CL | 443.25 | -1.293 | 0.154 | -8.38 | 5.26E-17 | 5.36E-15 | 14.330 | Downregulated |
| C10orf32-ASMT | 288.44 | -1.586 | 0.190 | -8.35 | 6.62E-17 | 6.57E-15 | 14.270 | Downregulated |
| GREM2 | 492.42 | -1.189 | 0.144 | -8.26 | 1.40E-16 | 1.36E-14 | 13.918 | Downregulated |
| OLFML1 | 516.66 | -1.165 | 0.143 | -8.16 | 3.25E-16 | 3.05E-14 | 13.565 | Downregulated |
| CYP4F12 | 494.23 | -1.355 | 0.169 | -8.00 | 1.28E-15 | 1.11E-13 | 13.025 | Downregulated |
| SLCO4C1 | 698.17 | -1.031 | 0.129 | -8.00 | 1.21E-15 | 1.05E-13 | 13.018 | Downregulated |
| LINC01089 | 108.66 | -2.385 | 0.301 | -7.93 | 2.14E-15 | 1.83E-13 | 12.958 | Downregulated |
| SRD5A2 | 224.50 | -1.427 | 0.198 | -7.22 | 5.14E-13 | 3.38E-11 | 10.568 | Downregulated |
| GOLGA8A | 155.08 | -1.744 | 0.242 | -7.20 | 6.16E-13 | 3.96E-11 | 10.547 | Downregulated |
| CAPN10-AS1 | 98.91 | -2.197 | 0.310 | -7.09 | 1.38E-12 | 8.46E-11 | 10.310 | Downregulated |
| LOC730101 | 422.30 | -1.267 | 0.178 | -7.14 | 9.61E-13 | 6.03E-11 | 10.298 | Downregulated |
| CSAD | 534.93 | -1.237 | 0.173 | -7.13 | 1.02E-12 | 6.33E-11 | 10.273 | Downregulated |
| IL17RB | 483.91 | -1.009 | 0.143 | -7.06 | 1.67E-12 | 9.98E-11 | 10.052 | Downregulated |
| ZNF292 | 553.80 | -1.021 | 0.146 | -7.00 | 2.55E-12 | 1.48E-10 | 9.883 | Downregulated |
| SNHG3 | 198.78 | -1.455 | 0.210 | -6.94 | 3.80E-12 | 2.14E-10 | 9.779 | Downregulated |
| HLA-G | 93.90 | -2.182 | 0.321 | -6.79 | 1.12E-11 | 5.86E-10 | 9.486 | Downregulated |
| PDK1 | 263.43 | -1.500 | 0.222 | -6.74 | 1.57E-11 | 8.15E-10 | 9.212 | Downregulated |
| PRLR | 128.13 | -1.826 | 0.277 | -6.60 | 4.17E-11 | 1.97E-09 | 8.895 | Downregulated |
| PMM1 | 292.16 | -1.167 | 0.177 | -6.59 | 4.41E-11 | 2.07E-09 | 8.761 | Downregulated |
| FBXL19-AS1 | 412.27 | -1.043 | 0.160 | -6.50 | 7.85E-11 | 3.50E-09 | 8.520 | Downregulated |
| LINC00528 | 53.95 | -2.750 | 0.433 | -6.35 | 2.13E-10 | 8.85E-09 | 8.510 | Downregulated |
| HERC5 | 225.50 | -1.263 | 0.197 | -6.41 | 1.50E-10 | 6.39E-09 | 8.291 | Downregulated |
| CYP2A6 | 282.17 | -1.135 | 0.177 | -6.40 | 1.55E-10 | 6.55E-09 | 8.262 | Downregulated |
| SLC27A5 | 299.94 | -1.302 | 0.209 | -6.22 | 4.98E-10 | 1.99E-08 | 7.811 | Downregulated |
| ALOX12P2 | 156.58 | -1.453 | 0.237 | -6.13 | 8.84E-10 | 3.40E-08 | 7.608 | Downregulated |
| LOC153910 | 89.40 | -1.918 | 0.318 | -6.04 | 1.54E-09 | 5.64E-08 | 7.498 | Downregulated |
| FETUB | 227.24 | -1.187 | 0.195 | -6.09 | 1.13E-09 | 4.29E-08 | 7.463 | Downregulated |
| SLC1A2 | 46.47 | -2.828 | 0.486 | -5.82 | 5.91E-09 | 1.96E-07 | 7.279 | Downregulated |
| CXCL10 | 54.20 | -2.431 | 0.417 | -5.83 | 5.51E-09 | 1.85E-07 | 7.159 | Downregulated |
| LINC01311 | 53.20 | -2.399 | 0.418 | -5.75 | 9.15E-09 | 2.93E-07 | 6.960 | Downregulated |
| PFKFB4 | 93.65 | -1.955 | 0.339 | -5.77 | 7.93E-09 | 2.57E-07 | 6.874 | Downregulated |
| PPFIA4 | 35.22 | -3.184 | 0.578 | -5.51 | 3.53E-08 | 1.00E-06 | 6.792 | Downregulated |
| NUSAP1 | 117.62 | -1.535 | 0.267 | -5.76 | 8.53E-09 | 2.75E-07 | 6.739 | Downregulated |
| AKR1C8P | 119.88 | -1.572 | 0.274 | -5.74 | 9.34E-09 | 2.98E-07 | 6.712 | Downregulated |
| SSUH2 | 69.69 | -2.129 | 0.376 | -5.67 | 1.45E-08 | 4.44E-07 | 6.700 | Downregulated |
| HNF1A-AS1 | 116.65 | -1.842 | 0.323 | -5.70 | 1.23E-08 | 3.83E-07 | 6.676 | Downregulated |
| PPP1R3E | 108.39 | -1.596 | 0.284 | -5.61 | 2.00E-08 | 5.91E-07 | 6.430 | Downregulated |
| GPR88 | 74.18 | -1.925 | 0.348 | -5.52 | 3.30E-08 | 9.45E-07 | 6.325 | Downregulated |
| INO80B | 254.97 | -1.121 | 0.202 | -5.54 | 3.05E-08 | 8.84E-07 | 6.157 | Downregulated |
| LOC729603 | 163.83 | -1.319 | 0.239 | -5.51 | 3.50E-08 | 9.94E-07 | 6.146 | Downregulated |
| ALDOC | 1169.63 | -1.422 | 0.259 | -5.49 | 4.09E-08 | 1.15E-06 | 6.108 | Downregulated |
| THUMPD3-AS1 | 150.84 | -1.331 | 0.243 | -5.47 | 4.50E-08 | 1.24E-06 | 6.054 | Downregulated |
| HMGCS2 | 4954.08 | -1.220 | 0.225 | -5.43 | 5.78E-08 | 1.58E-06 | 5.929 | Downregulated |
| FAM151A | 189.52 | -1.195 | 0.222 | -5.37 | 7.77E-08 | 2.04E-06 | 5.815 | Downregulated |
| DSCAML1 | 27.49 | -3.325 | 0.682 | -4.88 | 1.08E-06 | 2.20E-05 | 5.723 | Downregulated |
| MMP3 | 52.20 | -2.127 | 0.410 | -5.19 | 2.15E-07 | 5.13E-06 | 5.701 | Downregulated |
| DCDC5 | 69.19 | -1.888 | 0.362 | -5.21 | 1.88E-07 | 4.56E-06 | 5.665 | Downregulated |
| APBB3 | 206.75 | -1.074 | 0.202 | -5.31 | 1.10E-07 | 2.77E-06 | 5.660 | Downregulated |
| ABCB11 | 161.83 | -1.251 | 0.237 | -5.27 | 1.36E-07 | 3.37E-06 | 5.614 | Downregulated |
| THRSP | 855.15 | -1.428 | 0.272 | -5.24 | 1.58E-07 | 3.83E-06 | 5.602 | Downregulated |
| KMT2E-AS1 | 57.20 | -1.995 | 0.389 | -5.13 | 2.89E-07 | 6.73E-06 | 5.544 | Downregulated |
| ZNF436-AS1 | 84.18 | -1.758 | 0.341 | -5.16 | 2.47E-07 | 5.87E-06 | 5.519 | Downregulated |
| ASIC3 | 61.94 | -1.917 | 0.374 | -5.13 | 2.88E-07 | 6.72E-06 | 5.517 | Downregulated |
| ADAMTS17 | 125.86 | -1.339 | 0.258 | -5.19 | 2.11E-07 | 5.06E-06 | 5.462 | Downregulated |
| CYP17A1 | 24.98 | -3.173 | 0.668 | -4.75 | 2.03E-06 | 3.80E-05 | 5.441 | Downregulated |
| HSD17B14 | 82.41 | -1.623 | 0.317 | -5.11 | 3.19E-07 | 7.37E-06 | 5.383 | Downregulated |
| NEB | 1239.74 | -2.058 | 0.413 | -4.98 | 6.29E-07 | 1.35E-05 | 5.286 | Downregulated |
| CYP2B7P | 99.89 | -1.493 | 0.294 | -5.07 | 3.98E-07 | 9.04E-06 | 5.260 | Downregulated |
| CDCA2 | 134.08 | -1.465 | 0.290 | -5.05 | 4.37E-07 | 9.85E-06 | 5.217 | Downregulated |
| NFYC-AS1 | 56.95 | -1.948 | 0.393 | -4.96 | 7.01E-07 | 1.49E-05 | 5.204 | Downregulated |
| FASN | 14540.09 | -1.386 | 0.275 | -5.04 | 4.59E-07 | 1.02E-05 | 5.179 | Downregulated |
| OXER1 | 271.70 | -1.085 | 0.214 | -5.07 | 3.98E-07 | 9.04E-06 | 5.159 | Downregulated |
| APOM | 185.53 | -1.079 | 0.214 | -5.05 | 4.38E-07 | 9.85E-06 | 5.122 | Downregulated |
| C12orf76 | 167.06 | -1.131 | 0.224 | -5.04 | 4.55E-07 | 1.02E-05 | 5.119 | Downregulated |
| BNIP3 | 4825.32 | -1.285 | 0.256 | -5.01 | 5.44E-07 | 1.19E-05 | 5.091 | Downregulated |
| RNPC3 | 146.34 | -1.199 | 0.244 | -4.92 | 8.66E-07 | 1.82E-05 | 4.890 | Downregulated |
| TRIM31 | 88.90 | -1.502 | 0.309 | -4.86 | 1.15E-06 | 2.33E-05 | 4.871 | Downregulated |
| TIGD1 | 54.94 | -1.922 | 0.402 | -4.78 | 1.80E-06 | 3.43E-05 | 4.861 | Downregulated |
| RTN1 | 151.83 | -1.173 | 0.242 | -4.85 | 1.21E-06 | 2.42E-05 | 4.762 | Downregulated |
| C8orf44-SGK3 | 27.98 | -4.422 | 1.527 | -2.90 | 3.78E-03 | 2.42E-02 | 4.709 | Downregulated |
| GSTA2 | 1666.15 | -1.183 | 0.246 | -4.82 | 1.46E-06 | 2.87E-05 | 4.694 | Downregulated |
| MYOM1 | 180.05 | -1.088 | 0.225 | -4.83 | 1.38E-06 | 2.72E-05 | 4.693 | Downregulated |
| NEAT1 | 6897.57 | -1.305 | 0.274 | -4.77 | 1.86E-06 | 3.53E-05 | 4.639 | Downregulated |
| SPATA21 | 26.23 | -2.703 | 0.617 | -4.38 | 1.19E-05 | 1.87E-04 | 4.605 | Downregulated |
| SNHG19 | 69.18 | -1.679 | 0.359 | -4.68 | 2.94E-06 | 5.32E-05 | 4.592 | Downregulated |
| AKR1D1 | 157.06 | -1.114 | 0.234 | -4.76 | 1.89E-06 | 3.60E-05 | 4.582 | Downregulated |
| SPRY1 | 82.41 | -1.484 | 0.318 | -4.67 | 3.03E-06 | 5.46E-05 | 4.514 | Downregulated |
| TENM1 | 132.85 | -1.207 | 0.257 | -4.69 | 2.69E-06 | 4.90E-05 | 4.476 | Downregulated |
| TSSK3 | 122.88 | -1.376 | 0.299 | -4.61 | 4.07E-06 | 7.08E-05 | 4.372 | Downregulated |
| FST | 180.02 | -1.097 | 0.238 | -4.60 | 4.16E-06 | 7.21E-05 | 4.285 | Downregulated |
| MSC | 58.68 | -1.713 | 0.389 | -4.41 | 1.04E-05 | 1.65E-04 | 4.153 | Downregulated |
| RAB37 | 81.66 | -1.400 | 0.318 | -4.39 | 1.11E-05 | 1.75E-04 | 4.010 | Downregulated |
| CBLN3 | 44.70 | -1.894 | 0.446 | -4.25 | 2.18E-05 | 3.16E-04 | 3.980 | Downregulated |
| DLG4 | 123.36 | -1.145 | 0.262 | -4.38 | 1.20E-05 | 1.87E-04 | 3.900 | Downregulated |
| ADH1B | 1312.88 | -1.942 | 0.466 | -4.17 | 3.03E-05 | 4.21E-04 | 3.895 | Downregulated |
| MDM4 | 125.35 | -1.112 | 0.256 | -4.34 | 1.41E-05 | 2.18E-04 | 3.827 | Downregulated |
| IGF1 | 94.64 | -1.256 | 0.292 | -4.30 | 1.74E-05 | 2.63E-04 | 3.794 | Downregulated |
| C1orf220 | 62.20 | -1.658 | 0.401 | -4.14 | 3.49E-05 | 4.77E-04 | 3.712 | Downregulated |
| LDLRAD1 | 33.97 | -2.016 | 0.502 | -4.01 | 6.00E-05 | 7.64E-04 | 3.712 | Downregulated |
| VWCE | 2164.32 | -1.211 | 0.285 | -4.25 | 2.16E-05 | 3.13E-04 | 3.708 | Downregulated |
| LINC00115 | 38.71 | -1.889 | 0.466 | -4.06 | 4.99E-05 | 6.54E-04 | 3.703 | Downregulated |
| ADH1C | 102.90 | -1.406 | 0.336 | -4.19 | 2.79E-05 | 3.91E-04 | 3.687 | Downregulated |
| CCDC183 | 138.59 | -1.085 | 0.255 | -4.26 | 2.05E-05 | 3.01E-04 | 3.685 | Downregulated |
| LOC644656 | 33.96 | -2.016 | 0.505 | -3.99 | 6.64E-05 | 8.33E-04 | 3.680 | Downregulated |
| MALAT1 | 1072.63 | -1.290 | 0.309 | -4.17 | 3.01E-05 | 4.19E-04 | 3.616 | Downregulated |
| LOC101928505 | 90.14 | -1.250 | 0.300 | -4.17 | 3.01E-05 | 4.19E-04 | 3.602 | Downregulated |
| NIPBL-AS1 | 94.64 | -1.220 | 0.292 | -4.17 | 3.03E-05 | 4.21E-04 | 3.589 | Downregulated |
| STX16-NPEPL1 | 158.33 | -1.184 | 0.284 | -4.17 | 3.08E-05 | 4.27E-04 | 3.572 | Downregulated |
| SLC23A1 | 118.61 | -1.090 | 0.263 | -4.15 | 3.34E-05 | 4.59E-04 | 3.512 | Downregulated |
| LINC01004 | 94.15 | -1.301 | 0.317 | -4.10 | 4.19E-05 | 5.62E-04 | 3.501 | Downregulated |
| FGF11 | 81.15 | -1.432 | 0.354 | -4.04 | 5.34E-05 | 6.92E-04 | 3.469 | Downregulated |
| CYP2D7 | 43.71 | -1.758 | 0.450 | -3.90 | 9.56E-05 | 1.14E-03 | 3.427 | Downregulated |
| TTLL7 | 105.12 | -1.149 | 0.283 | -4.06 | 4.85E-05 | 6.37E-04 | 3.396 | Downregulated |
| HIST1H4E | 49.20 | -1.677 | 0.433 | -3.87 | 1.07E-04 | 1.26E-03 | 3.350 | Downregulated |
| LPPR1 | 41.21 | -1.743 | 0.456 | -3.82 | 1.32E-04 | 1.51E-03 | 3.315 | Downregulated |
| ATP1A2 | 66.69 | -1.440 | 0.370 | -3.89 | 9.95E-05 | 1.19E-03 | 3.261 | Downregulated |
| DKFZP434I0714 | 50.44 | -1.531 | 0.400 | -3.83 | 1.30E-04 | 1.49E-03 | 3.215 | Downregulated |
| LOC646471 | 119.62 | -1.122 | 0.285 | -3.94 | 8.06E-05 | 9.82E-04 | 3.210 | Downregulated |
| SPOCK1 | 123.35 | -1.025 | 0.259 | -3.96 | 7.52E-05 | 9.25E-04 | 3.202 | Downregulated |
| ARNT2 | 29.47 | -1.972 | 0.545 | -3.62 | 2.93E-04 | 3.00E-03 | 3.202 | Downregulated |
| MIR17HG | 27.22 | -1.986 | 0.557 | -3.57 | 3.58E-04 | 3.56E-03 | 3.153 | Downregulated |
| TENM2 | 99.38 | -1.113 | 0.291 | -3.82 | 1.34E-04 | 1.53E-03 | 3.028 | Downregulated |
| ASPA | 43.20 | -1.599 | 0.441 | -3.63 | 2.85E-04 | 2.93E-03 | 2.996 | Downregulated |
| APOF | 116.37 | -1.064 | 0.280 | -3.80 | 1.43E-04 | 1.62E-03 | 2.987 | Downregulated |
| GPRIN3 | 35.21 | -1.713 | 0.481 | -3.56 | 3.69E-04 | 3.64E-03 | 2.980 | Downregulated |
| UNC13D | 48.46 | -1.607 | 0.446 | -3.61 | 3.11E-04 | 3.15E-03 | 2.973 | Downregulated |
| MIR6723 | 39.21 | -1.650 | 0.467 | -3.53 | 4.14E-04 | 4.02E-03 | 2.909 | Downregulated |
| NDRG1 | 11395.50 | -1.430 | 0.398 | -3.59 | 3.26E-04 | 3.28E-03 | 2.866 | Downregulated |
| HEXA-AS1 | 28.47 | -1.835 | 0.544 | -3.37 | 7.46E-04 | 6.51E-03 | 2.854 | Downregulated |
| HSD17B7P2 | 40.46 | -1.564 | 0.449 | -3.49 | 4.88E-04 | 4.60E-03 | 2.812 | Downregulated |
| STX1B | 104.12 | -1.018 | 0.276 | -3.68 | 2.29E-04 | 2.45E-03 | 2.803 | Downregulated |
| TRPV3 | 74.92 | -1.202 | 0.335 | -3.59 | 3.27E-04 | 3.29E-03 | 2.759 | Downregulated |
| HK2 | 91.40 | -2.008 | 0.641 | -3.13 | 1.73E-03 | 1.30E-02 | 2.756 | Downregulated |
| TLR9 | 37.21 | -1.601 | 0.471 | -3.40 | 6.73E-04 | 5.97E-03 | 2.740 | Downregulated |
| LINC01558 | 25.97 | -1.819 | 0.563 | -3.23 | 1.22E-03 | 9.77E-03 | 2.711 | Downregulated |
| CA9 | 377.83 | -2.439 | 1.037 | -2.35 | 1.86E-02 | 8.17E-02 | 2.670 | Downregulated |
| RDH16 | 46.20 | -1.435 | 0.421 | -3.41 | 6.54E-04 | 5.83E-03 | 2.655 | Downregulated |
| PNCK | 41.20 | -2.356 | 0.947 | -2.49 | 1.28E-02 | 6.20E-02 | 2.648 | Downregulated |
| SNHG21 | 31.73 | -1.760 | 0.553 | -3.18 | 1.46E-03 | 1.13E-02 | 2.625 | Downregulated |
| SNHG20 | 60.68 | -1.279 | 0.371 | -3.44 | 5.72E-04 | 5.21E-03 | 2.617 | Downregulated |
| PCED1B | 66.68 | -1.202 | 0.349 | -3.45 | 5.67E-04 | 5.19E-03 | 2.582 | Downregulated |
| SLC25A27 | 102.14 | -1.121 | 0.323 | -3.47 | 5.18E-04 | 4.83E-03 | 2.573 | Downregulated |
| SPARCL1 | 32.71 | -1.632 | 0.509 | -3.20 | 1.35E-03 | 1.06E-02 | 2.563 | Downregulated |
| APOC4-APOC2 | 29.47 | -1.687 | 0.537 | -3.14 | 1.68E-03 | 1.27E-02 | 2.539 | Downregulated |
| MSC-AS1 | 30.71 | -1.635 | 0.519 | -3.15 | 1.65E-03 | 1.25E-02 | 2.510 | Downregulated |
| LOC100128398 | 67.43 | -1.200 | 0.356 | -3.37 | 7.43E-04 | 6.49E-03 | 2.495 | Downregulated |
| LOC171391 | 93.64 | -1.003 | 0.292 | -3.43 | 6.03E-04 | 5.45E-03 | 2.476 | Downregulated |
| DNAJC3-AS1 | 56.44 | -1.249 | 0.375 | -3.33 | 8.66E-04 | 7.32E-03 | 2.474 | Downregulated |
| C15orf48 | 120.10 | -1.723 | 0.583 | -2.95 | 3.13E-03 | 2.07E-02 | 2.409 | Downregulated |
| SNHG10 | 39.46 | -1.468 | 0.467 | -3.15 | 1.66E-03 | 1.25E-02 | 2.403 | Downregulated |
| COL16A1 | 109.62 | -1.007 | 0.300 | -3.36 | 7.91E-04 | 6.79E-03 | 2.391 | Downregulated |
| TMCO6 | 83.40 | -1.029 | 0.309 | -3.33 | 8.57E-04 | 7.26E-03 | 2.374 | Downregulated |
| EFNA1 | 1404.95 | -1.323 | 0.415 | -3.19 | 1.43E-03 | 1.10E-02 | 2.362 | Downregulated |
| CAPN3 | 106.14 | -1.013 | 0.304 | -3.33 | 8.73E-04 | 7.36E-03 | 2.361 | Downregulated |
| PTPRH | 67.41 | -1.199 | 0.372 | -3.22 | 1.27E-03 | 1.01E-02 | 2.329 | Downregulated |
| ALOX15B | 52.94 | -1.244 | 0.389 | -3.20 | 1.39E-03 | 1.08E-02 | 2.326 | Downregulated |
| RGAG1 | 33.96 | -1.477 | 0.487 | -3.03 | 2.43E-03 | 1.70E-02 | 2.305 | Downregulated |
| EXPH5 | 63.68 | -1.132 | 0.351 | -3.22 | 1.28E-03 | 1.01E-02 | 2.293 | Downregulated |
| CEMP1 | 107.87 | -1.032 | 0.317 | -3.26 | 1.11E-03 | 8.97E-03 | 2.293 | Downregulated |
| DIRAS2 | 44.95 | -1.302 | 0.417 | -3.13 | 1.77E-03 | 1.32E-02 | 2.287 | Downregulated |
| LOC100506127 | 54.95 | -1.288 | 0.412 | -3.12 | 1.78E-03 | 1.32E-02 | 2.278 | Downregulated |
| COLCA1 | 41.96 | -1.410 | 0.463 | -3.04 | 2.35E-03 | 1.65E-02 | 2.273 | Downregulated |
| SLC6A13 | 37.71 | -1.426 | 0.483 | -2.96 | 3.12E-03 | 2.07E-02 | 2.207 | Downregulated |
| IGSF1 | 54.93 | -1.163 | 0.379 | -3.07 | 2.14E-03 | 1.53E-02 | 2.155 | Downregulated |
| NRN1 | 36.21 | -1.395 | 0.481 | -2.90 | 3.75E-03 | 2.40E-02 | 2.137 | Downregulated |
| LHX4-AS1 | 27.48 | -1.623 | 0.633 | -2.56 | 1.03E-02 | 5.26E-02 | 2.066 | Downregulated |
| GP1BA | 44.20 | -1.229 | 0.425 | -2.89 | 3.85E-03 | 2.46E-02 | 2.025 | Downregulated |
| UTS2B | 38.21 | -1.311 | 0.466 | -2.82 | 4.86E-03 | 2.95E-02 | 2.015 | Downregulated |
| LBHD1 | 69.93 | -1.057 | 0.360 | -2.94 | 3.31E-03 | 2.17E-02 | 1.971 | Downregulated |
| NEURL2 | 33.21 | -1.325 | 0.485 | -2.73 | 6.32E-03 | 3.60E-02 | 1.960 | Downregulated |
| LOC440434 | 70.92 | -1.041 | 0.357 | -2.91 | 3.57E-03 | 2.31E-02 | 1.939 | Downregulated |
| ZNF674-AS1 | 35.21 | -1.289 | 0.472 | -2.73 | 6.37E-03 | 3.61E-02 | 1.934 | Downregulated |
| C9orf173-AS1 | 37.46 | -1.365 | 0.515 | -2.65 | 8.00E-03 | 4.34E-02 | 1.929 | Downregulated |
| SLC22A25 | 35.46 | -1.303 | 0.484 | -2.70 | 7.04E-03 | 3.91E-02 | 1.918 | Downregulated |
| CFAP57 | 26.72 | -1.430 | 0.559 | -2.56 | 1.05E-02 | 5.32E-02 | 1.915 | Downregulated |
| DUSP5P1 | 49.45 | -1.136 | 0.403 | -2.82 | 4.78E-03 | 2.90E-02 | 1.911 | Downregulated |
| APLN | 36.45 | -1.263 | 0.470 | -2.69 | 7.25E-03 | 4.00E-02 | 1.884 | Downregulated |
| PAXIP1-AS1 | 65.43 | -1.011 | 0.353 | -2.86 | 4.20E-03 | 2.63E-02 | 1.876 | Downregulated |
| GUCY2D | 33.96 | -1.266 | 0.480 | -2.64 | 8.38E-03 | 4.50E-02 | 1.848 | Downregulated |
| EGFR-AS1 | 182.01 | -1.270 | 0.487 | -2.61 | 9.12E-03 | 4.79E-02 | 1.831 | Downregulated |
| OXT | 58.18 | -1.021 | 0.365 | -2.80 | 5.18E-03 | 3.08E-02 | 1.824 | Downregulated |
| SKAP1 | 34.21 | -1.280 | 0.499 | -2.57 | 1.03E-02 | 5.24E-02 | 1.811 | Downregulated |
| NEFL | 31.21 | -1.308 | 0.516 | -2.53 | 1.13E-02 | 5.64E-02 | 1.809 | Downregulated |
| AQP7 | 50.19 | -1.101 | 0.405 | -2.72 | 6.61E-03 | 3.72E-02 | 1.804 | Downregulated |
| IL1RN | 53.93 | -1.033 | 0.377 | -2.74 | 6.11E-03 | 3.52E-02 | 1.783 | Downregulated |
| ANXA10 | 35.71 | -1.220 | 0.473 | -2.58 | 9.87E-03 | 5.09E-02 | 1.778 | Downregulated |
| TMEM44-AS1 | 55.93 | -1.022 | 0.376 | -2.72 | 6.53E-03 | 3.68E-02 | 1.761 | Downregulated |
| B4GALNT1 | 42.20 | -1.133 | 0.433 | -2.62 | 8.86E-03 | 4.69E-02 | 1.746 | Downregulated |
| FZD7 | 54.44 | -1.023 | 0.379 | -2.70 | 6.91E-03 | 3.85E-02 | 1.746 | Downregulated |
| SCML4 | 39.95 | -1.140 | 0.437 | -2.61 | 9.13E-03 | 4.80E-02 | 1.743 | Downregulated |
| RNF144A | 54.43 | -1.053 | 0.395 | -2.67 | 7.63E-03 | 4.18E-02 | 1.735 | Downregulated |
| CNTNAP2 | 28.22 | -1.280 | 0.526 | -2.43 | 1.49E-02 | 6.96E-02 | 1.726 | Downregulated |
| EGLN3 | 1761.91 | -1.109 | 0.426 | -2.60 | 9.19E-03 | 4.82E-02 | 1.722 | Downregulated |
| LOC100289230 | 27.47 | -1.288 | 0.542 | -2.38 | 1.74E-02 | 7.74E-02 | 1.701 | Downregulated |
| SPAG4 | 227.70 | -1.096 | 0.426 | -2.57 | 1.00E-02 | 5.15E-02 | 1.691 | Downregulated |
| STXBP6 | 38.45 | -1.234 | 0.508 | -2.43 | 1.52E-02 | 7.03E-02 | 1.689 | Downregulated |
| KIAA1875 | 34.21 | -1.180 | 0.477 | -2.48 | 1.33E-02 | 6.37E-02 | 1.680 | Downregulated |
| ZNF471 | 36.20 | -1.155 | 0.465 | -2.48 | 1.30E-02 | 6.25E-02 | 1.668 | Downregulated |
| NDUFA4L2 | 2936.87 | -1.091 | 0.432 | -2.53 | 1.15E-02 | 5.72E-02 | 1.654 | Downregulated |
| CIT | 49.70 | -1.047 | 0.411 | -2.54 | 1.09E-02 | 5.50E-02 | 1.638 | Downregulated |
| LOC100272217 | 32.46 | -1.173 | 0.490 | -2.39 | 1.68E-02 | 7.54E-02 | 1.623 | Downregulated |
| TREX1 | 46.20 | -1.062 | 0.424 | -2.50 | 1.24E-02 | 6.03E-02 | 1.617 | Downregulated |
| HERC2P7 | 37.96 | -1.118 | 0.466 | -2.40 | 1.65E-02 | 7.46E-02 | 1.587 | Downregulated |
| C8orf46 | 42.70 | -1.041 | 0.425 | -2.45 | 1.43E-02 | 6.74E-02 | 1.567 | Downregulated |
| LINC00641 | 37.22 | -1.166 | 0.506 | -2.30 | 2.12E-02 | 9.00E-02 | 1.566 | Downregulated |
| REC8 | 29.97 | -1.168 | 0.512 | -2.28 | 2.26E-02 | 9.40E-02 | 1.556 | Downregulated |
| STAG3L3 | 34.21 | -1.131 | 0.488 | -2.32 | 2.06E-02 | 8.78E-02 | 1.548 | Downregulated |
| SLC25A21-AS1 | 40.70 | -1.056 | 0.442 | -2.39 | 1.68E-02 | 7.55E-02 | 1.541 | Downregulated |
| ECM1 | 37.95 | -1.074 | 0.454 | -2.37 | 1.80E-02 | 7.96E-02 | 1.537 | Downregulated |
| MEX3B | 32.96 | -1.102 | 0.481 | -2.29 | 2.20E-02 | 9.24E-02 | 1.512 | Downregulated |
| PRRT2 | 36.46 | -1.077 | 0.478 | -2.25 | 2.42E-02 | 9.90E-02 | 1.473 | Downregulated |
| HIST4H4 | 34.71 | -1.065 | 0.471 | -2.26 | 2.36E-02 | 9.69E-02 | 1.471 | Downregulated |

**S2 Table　Antibodies used in this study**

| **Abbreviated name** | **Official name** | **Company name** | **Catalog No.** | **Diluted concentration** |
| --- | --- | --- | --- | --- |
| Rabbit | Peroxidase AffiniPure Goat Anti-Rabbit IgG (H+L) | Jackson Immuno Research | 111-035-003 | 1:10,000 |
| Mouse | Goat anti-Mouse IgG (H+L) Secondary Antibody, HRP | Thermo Fisher Scientific | 31430 | 1:10,000 |
| HA | Purified anti-HA.11 Epitope Tag Antibody | BioLegend | 901501 | 1:1,000 |
| GAPDH | Anti GAPDH, Monoclonal Antibody, Peroxidase Conjugated | Fujifilm | 015-25473 | 1:10,000 |
| N4BP1 | N4BP1 Antibody | Bethyl | A304-628A-T | 1:1,000 |
| ZCCHC10 | Anti-ZCCHC10 antibody produced in rabbit | Sigma | HPA038944 | 1:500 |
| HNRNPC | Anti-hnRNP C1 + C2/HNRNPC 抗体 [EP3034Y] | abcam | ab75822 | 1:1,000 |
| KHNYN | KHNYN Antibody | Fujifilm | NBP2-17041 | 1:1,000 |
| ZC3H12B | ZC3H12B antibody | Gene Tex | GTX85196 | 1:1,000 |
